# Toroidal topology of population activity in grid cells

**DOI:** 10.1101/2021.02.25.432776

**Authors:** Richard J. Gardner, Erik Hermansen, Marius Pachitariu, Yoram Burak, Nils A. Baas, Benjamin A. Dunn, May-Britt Moser, Edvard I. Moser

## Abstract

The medial entorhinal cortex (MEC) is part of a neural system for mapping a subject’s position within a physical environment^1,2^. Grid cells, a key component of this system, fire in a characteristic hexagonal pattern of locations^3^, and are organized in modules^4,5^ which collectively form a population code for the animal’s allocentric position^1,6–8^. The invariance of the correlation structure of this population code across environments^9,10^ and behavioural states^11,12^, independently of specific sensory inputs, has pointed to intrinsic, recurrently connected continuous attractor networks (CANs) as a possible substrate of the grid pattern^1,2,13–16^. However, whether grid cell networks show continuous attractor dynamics, and how they interface with inputs from the environment, has remained elusive due to the small samples of cells obtained to date. Here we show, with simultaneous recordings from many hundreds of grid cells, and subsequent topological data analysis, that the joint activity of grid cells from an individual module resides on a toroidal manifold, as expected in a two-dimensional CAN. Positions on the torus correspond to the moving animal’s position in the environment. Individual cells are preferentially active at singular positions on the torus. Their positions are maintained, with minimal distortion, between environments and from wakefulness to sleep, as predicted by CAN models for grid cells but not by alternative feed-forward models where grid patterns are created from external inputs by Hebbian plasticity^17–22^. This demonstration of network dynamics on a toroidal manifold provides the first population-level visualization of CAN dynamics in grid cells.

## Main

The idea of a CAN has become one of the most influential concepts in theoretical systems neuroscience^23–25^. In these networks, recurrent synaptic connectivity constrains the joint activity of cells to a restricted, but continuous repertoire of possible coactivation patterns. CANs have been introduced to account for a wide range of brain functions that operate on continuous scales, spanning from visual orientation tuning^24^ to neural operations underlying motor control, decision making and action selection^26–28^, and certain forms of memory^29–33^. Few brain systems, however, are more suitable for analysis of CAN dynamics than spatial mapping circuits^2,34^, due to the continuous, low-dimensional nature of space, and the availability and interpretability of data from such circuits. In the mammalian hippocampal and parahippocampal cortices, head direction cells^35^ encode orientation whereas place cells^36^ and grid cells^3^ encode position. CAN models suggest that these cells operate on one-dimensional^37–39^ or two-dimensional^1,13–16,40–42^ continua that for head direction cells and grid cells are periodic. Head direction cells are arranged on a ring according to firing direction^37–39^, and grid cells on a torus according to firing location^1,13–16^. Because, in these models, inhibitory connections extend further across the neural continuum than excitatory connections, the CAN’s activity stabilizes as a localized bump on the ring or the torus. During navigation, this bump can be translated along the network continuum by speed and direction inputs, in accordance with the animal’s changing orientation or position in the external spatial environment.

While CAN models provide a coherent theoretical framework for the dynamics of spatially tuned cells, evidence for their existence in mammals remains indirect. The most direct evidence has recently appeared in flies, where CAN-like dynamics has been visualized in a ring of serially connected orientation-tuned cells of the central complex^43–45^. In rodents, head direction cedls^46–49^ and grid cells^9–12^ maintain fixed correlation structures, and in grid cells these are confined to modules^4,5^, as predicted by CAN models^1,13^. Yet, the small sample sizes of past studies have impeded experimental access to the full topology of the population activity. In rodent head direction cells, cell samples of a few dozen have been sufficient to demonstrate that network activity traverses a ring, in correspondence with the animal’s head direction^50–52^. For grid cells and place cells, however, which operate in two dimensions, the number of possible locations in state space is too large for the manifold to be uncovered with conventional sample sizes. Here we address this challenge for grid cells by taking advantage of recently developed high-site-count Neuropixels silicon probes^53,54^, which allow spike activity to be recorded simultaneously from thousands of cells in freely moving rats. Equipped with such probes, we set out to determine whether, as predicted by two-dimensional CAN models, the activity of the grid cell population in MEC resides on a toroidal manifold and whether, as predicted from the same models, this arrangement transcends behavioural tasks and states, decoupled from the animal’s position in physical space.

### Visualization of toroidal manifold

We recorded extracellular spikes of a total of 7,671 single units in layers II and III of the MEC-parasubiculum region in freely moving rats with unilateral or bilateral implants (total of 4 recordings, in 2 rats with bilateral single-shank probes and 1 rat with a unilateral 4-shank probe, from 546 to 2,460 cells per recording; Extended Data Fig. 1, Online methods). During recordings, the animals were engaged in foraging behaviour in a square recording enclosure or on an elevated track, or they slept in a small resting box. We employed a novel approach for assigning module identity to co-recorded grid cells, by clustering the periodic spatial firing patterns of the cells in the open-field arena (see Online methods). Six grid modules were clearly distinguished in this analysis (4 recording sessions, from 140 to 483 grid cells per session; Extended Data Fig. 2a-d). Each grid module cluster contained a mixture of nondirectional (pure) grid cells and conjunctive grid × direction cells^55^. We limited our analyses to the subset of pure grid cells because (i) the expected toroidal topology might be distorted by additional directional modulation, and (ii) detection of topology in conjunctive cells may require a larger number of cells than recorded here^56^.

To visually inspect the population activity structure of grid cells for signatures of toroidal topology, we constructed a 3-D embedding of the *N*-dimensional population activity of a module of *N* = 149 grid cells (Fig. 1a). For this, we applied a two-stage dimensionality reduction procedure on the matrix of firing rates. First, we conducted a principal component analysis (PCA) and kept the first six principal components, which gave the lowest-dimensional Euclidean embedding of the population activity that faithfully preserved the grid structure (Extended Data Fig. 3a). These first six principal components explained a particularly large fraction of the population activity (Extended Data Fig. 3b, c), and their values varied with grid-like periodicity as a function of the animal’s location in the open field (Extended data Fig. 3d). We next applied a second, nonlinear, dimensionality-reduction method (Uniform Manifold Approximation and Projection, UMAP), in which we further reduced the six-dimensional data to a 3-D visualization. This visualization revealed a striking torus-like structure (Fig. 1b; Supplementary Movie 1). Movement of the animal in the open field (OF) was accompanied by similarly continuous movement of the population activity across the toroidal manifold (Fig. 1b). When the activity of individual cells was plotted with reference to the 3-D population representation, spikes for each cell were localized within a single patch of the population state space (Fig. 1c). The offsets between individual cells’ firing locations in the arena corresponded with the cells’ relative firing locations in the toroidal state space.

**Figure 1:**
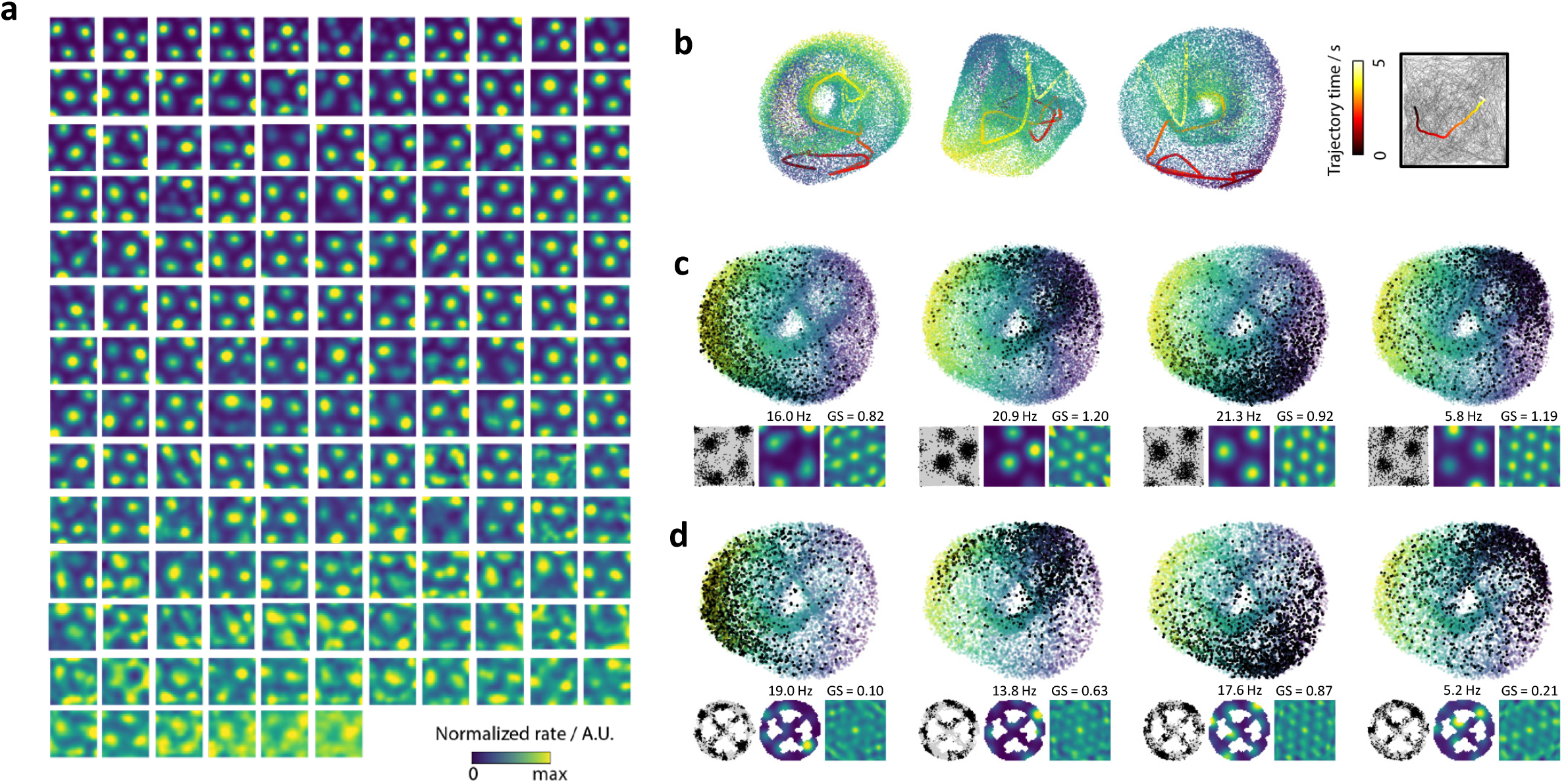
Signatures of toroidal structure in the activity of a module of grid cells. **a.** Firing rates of 149 grid cells co-recorded from the same module when a rat was foraging in an open-field arena (rat ‘R’ day 1, module 2; Extended Data Fig. 2). Firing rate is colour-coded as a function of the rat’s position in the arena. Colour coding is indicated by scale bar. Each square shows the activity map for one cell. Maximum rates range from 0.2 to 35.0 Hz. Cells are ranked in order of spatial information content (highest spatial information at top left, lowest at bottom right). Time periods when the animal’s speed was below 2.5 cm/s are excluded from all plots in this figure. **b.** Nonlinear dimensionality reduction reveals torus-like structure in population activity of a single grid module (same 149 cells as in **a**). PCA was performed to identify the six principal components with the highest variance explained (Extended Data Fig. 3). These components were further reduced to three dimensions by UMAP, yielding a 3-D nonlinear embedding, which is plotted here (3 different views of the same point cloud). Each dot of each point cloud represents the population activity state at a single time point. Dots are coloured by the value of the first principal component, to aid 3-D visualization. The bold coloured line shows a 5-s example trajectory, demonstrating smooth movement over the toroidal manifold. The corresponding behavioural trajectory of the animal in the open field is shown to the right. In all plots, colour indicates elapsed time (scale bar). **c.** Toroidal positions of spikes from four grid cells belonging to the module shown in **a**. Each panel shows the same 3-D point cloud of population activity states as in **b**, together with black dots which indicate the population state at times when the cell fired. Insets below each point cloud show (left) the 2-D firing locations of the cell in the open field (black dots superimposed on trajectory in grey), (middle) the colour-coded firing rate map of the cell in the open field, and (right) a similarly colour-coded autocorrelogram of the rate map. Colour scales of rate maps range from zero (dark blue) to the maximum value of the rate map for each cell. Colour scales for autocorrelograms range from −1 to +1. Text above rate maps indicates the maximum firing rate of each cell. Text above autocorrelograms indicates the grid score of the autocorrelogram. **d.** Same as **c**, plotted for the same four cells in a different behavioural session when the animal ran on an elevated, wheel-shaped track (“wagon-wheel track”).

### Quantification of toroidal topology

While the UMAP projection allowed a toroidal point cloud to be visualized, the method is reliant on optimization and sensitive to initial conditions^57^, and does not lend itself to straightforward quantification of the topology of the state space, nor does it easily permit comparison of representations across experiments. We therefore turned to the framework of persistent cohomology^58^, a toolset from topological data analysis in which the structure of neural data can be classified by identifying holes of varying dimensionality in topological spaces assigned to point clouds of the cells’ firing rates^50,51,59^ (Fig. 2). In applying this toolset, we replace each point of the point cloud by a ball of common radius. The union of balls results in a topological space where the number of holes of different dimensions can be counted. By increasing this radius from zero until all the balls intersect, we observe how each hole first appears at a given radius and subsequently disappears at a larger radius (see Fig. 2b). The range of radii for which each hole can be detected is referred to as the lifetime of the hole and is represented by the length of a bar. The totality of bars, for all holes, is referred to as the barcode. For ring topology, the barcode must display one bar of substantial length in dimension 0 (0-D hole; a single component connecting all points) and one in dimension one (1-D hole; a circle) (Fig. 2a), as shown for head direction cells^50–52^. For a torus, the barcode must display four bars of substantial length: one in dimension 0, two in dimension 1, and one in dimension 2 (2-D hole; a cavity).

**Figure 2:**
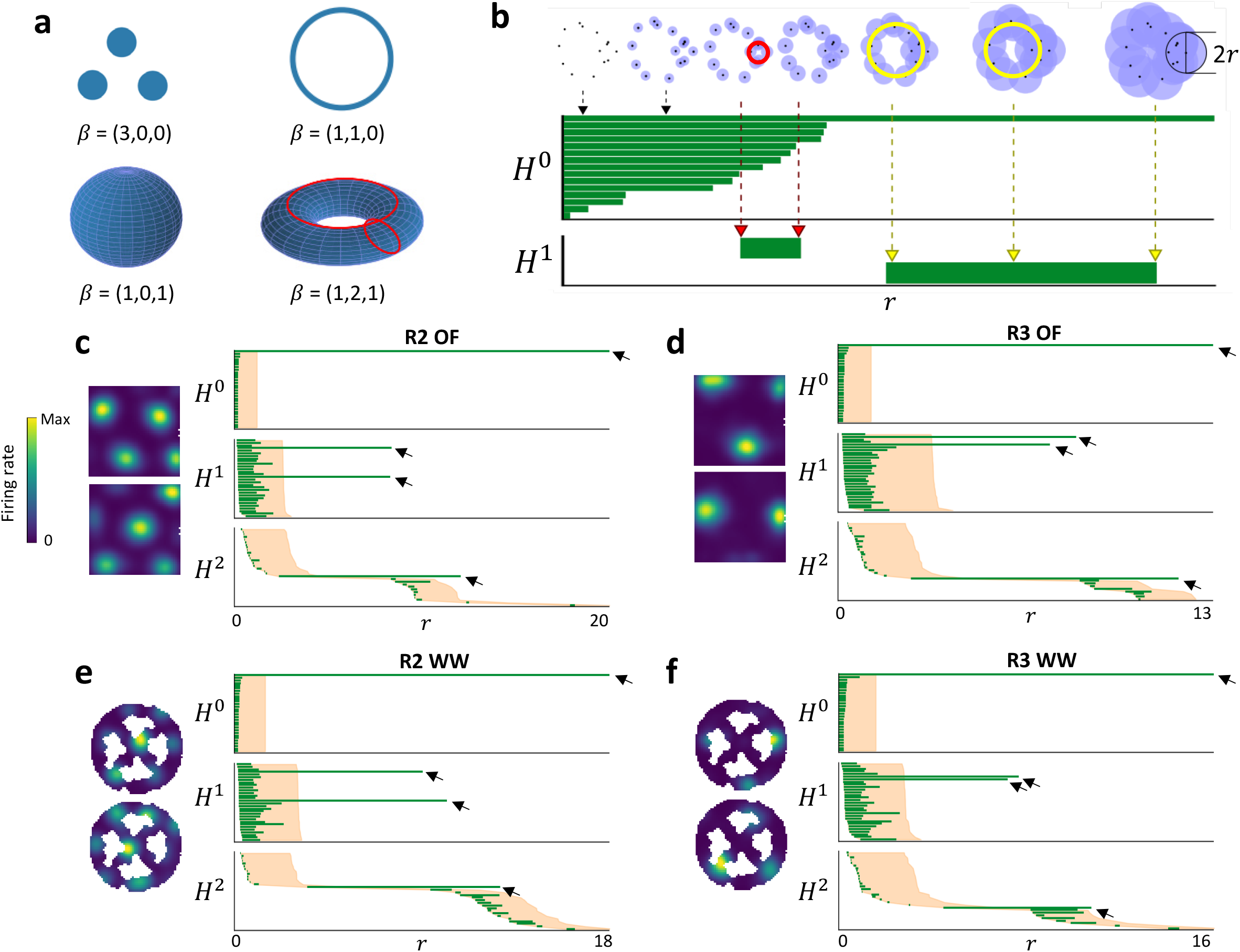
Persistent cohomology reveals toroidal structure in point clouds of grid-cell population activity. **a.** Cohomology can help differentiate topological spaces such as the union of three discs (upper left), a circle (upper right), a sphere (lower left) and a torus (lower right) by counting the number of topological holes in different dimensions. The Betti number of the space (*β*) is here given as a list of the number of holes in the first three dimensions. A disc has a 0-D hole (a connected component); a circle additionally has a 1-D hole; a (hollow) sphere is a connected component and has a 2-D hole (a cavity); a torus is a connected component with two 1-D holes (illustrated with red circles) and one 2-D hole (a cavity in the interior of the torus). **b.** Persistent cohomology tracks the lifetime of topological holes in spaces associated with point clouds. Top: The radius of balls centred at each data point in the point cloud is continuously increased (left to right). The union of the balls forms a space with possible holes. The lifetime of a hole during expansion of the radius is defined as the radial interval from when the hole first appears until it is filled in. Note the short lifetime of the hole marked with a red circle and the long lifetime of the hole indicated with a yellow circle. Second and third row: The lifetime of each hole of dimension zero (*H°*) and one (*H^1^)* in the example in the top row is indicated by the length of a bar (in green) in the *barcode* diagram. Two 1-D holes are detected: the first bar, corresponding to the red hole in the top row, is short and regarded as noise, while the second, corresponding to the yellow hole, is substantially longer and captures the prominent topology of the point cloud. **c., d.** Barcodes indicate toroidal topology of grid-cell population activity during open field (OF) foraging (**c**, module R2; **d**, module R3). Left of each panel: colour-coded firing rate maps for two representative grid cells in each module. Colour coding indicated by scale bar. Note difference in grid scale between R2 and R3. Right of each panel: Barcodes as in **b**. Arrows point to the four most prominent bars. These are substantially longer than a distribution of shuffled data from the same recording. The longest lifetime for each dimension in analyses of 1000 shuffles of the data is indicated by the width of the shaded orange region, with the left edge starting at the same value as each of the original bars. Note that in the original data, there is one long bar in dimension 0, two in dimension 1 and one in dimension 2, indicating Betti numbers equal to those of a torus. Only the 30 longest bars are shown for each dimension. **e., f.** Toroidal structure in an environment of higher topological complexity than the open field: the “wagon wheel” track (WW), as previously seen in Fig. 1d. Data are from the same grid modules as in **c** and **d**. Although grid cells had lower periodicity in WW than OF (Extended Data Fig. 2f), barcode diagrams are similar to those of open field data, suggesting preserved toroidal structure during distortions of the grid pattern.

Persistent cohomology analyses allowed us to classify the shape of the six-dimensional representation that serves as an intermediate step in UMAP (Extended Data Fig. 3a5). We constructed barcodes for each of the 6 individual modules of grid cells recorded in the open-field arena (3 modules from rat ‘R’, 2 from rat ‘Q’ and 1 from rat ‘S’, henceforth named R1, R2, R3, Q1, Q2 and S1). The barcodes showed clear indications of toroidal characteristics. For all 6 modules, we detected four long-lived bars representing a single 0-D hole, two 1-D holes and a 2-D hole. Their lifetimes were much longer than any other feature (bar) of the point cloud and significantly longer than the lifetime of any bar obtained in 1,000 shuffled versions of the data where the spike trains were shifted individually and randomly in time (Fig. 2c, d; Extended Data Fig. 5a; *P* < 0.001). The findings suggest that network dynamics during OF foraging resides on a low-dimensional manifold with the same barcode as a torus. We noted the appearance of additional short bars in the barcodes for all modules, but these are expected for toroidal point clouds^56^, as we confirmed with simulated data from several CAN models^15,16^ and point clouds sampled from idealized tori, in each case exhibiting similar features (see Extended Data Fig. 6).

### Persistence of toroidal topology in environments with reduced grid symmetry

A central feature of CAN models is that neural activity resides faithfully on a low-dimensional manifold that is mapped onto the physical environment by external sensory inputs, along with a velocity integration mechanism. The appearance of a torus in the point cloud, and the mapping of individual grid cells’ activity onto the torus (Fig. 1c), is consistent with a relationship between position in 2D physical space and in the dimensionality-reduced neural state space. However, in many environments, this relationship may not be isometric, since the grid pattern is distorted by geometrical features of the environment, such as walls and corners^4,60–62^ or discrete landmarks and reward locations^63,64^. We thus asked whether such geometric features could similarly distort the toroidal organization of network activity in the point cloud. To address this question, the animals were tested on an elevated running track shaped like a wagon wheel with four radial spokes (“wagon-wheel track” (WW), Fig. 1d and 2e, f). Rewards were given at eight fixed locations with salient visual features. Spatial autocorrelation analyses confirmed that the strict periodicity of the grid pattern was compromised in this task, even when differences in position coverage in the two environments were controlled for by “masking” the OF position coordinates to give similar spatial coverage between the two conditions (Extended Data Fig. 2 e, f; Online methods). In all modules, grid scores in the “masked” OF condition were higher than in WW (grid score mean ± S.E.M. across all cells: OF: 0.677 ± 0.017, WW: 0.360 ± 0.017, *N* = 618 cells, *P-* values for the 6 modules ranged from 1.26 × 10^-14^ to 0.03, Wilcoxon sign-rank test). Despite these distortions of the grid pattern in individual cells, toroidal tuning was maintained in the transformed population activity (Fig. 1d). The persistent cohomology analysis continued to identify one 0-D hole, two 1-D holes and one 2-D hole with lifetimes substantially exceeding those of a distribution of shuffled data (Fig. 2d and Extended Data Fig. 5a). Furthermore, grid cells in each module were individually tuned to the toroidal surface. This was demonstrated in a toroidal parametrization of the population activity (mapping the activity at each time point to a point on the torus) where we calculated angular coordinates from each of the two 1-D holes (‘cohomological decoding’, Online Methods; Extended Data Fig. 4). The two angular coordinates defined directions which intersected at 60 degrees, defining a rhombus, identifiable as a twisted torus. Consistent with CAN models, the vast majority of cells were tuned to a single location on the torus in each module and across environments, independent of geometry and local landmarks (Fig. 3b, Extended Data Fig. 9).

**Figure 3:**
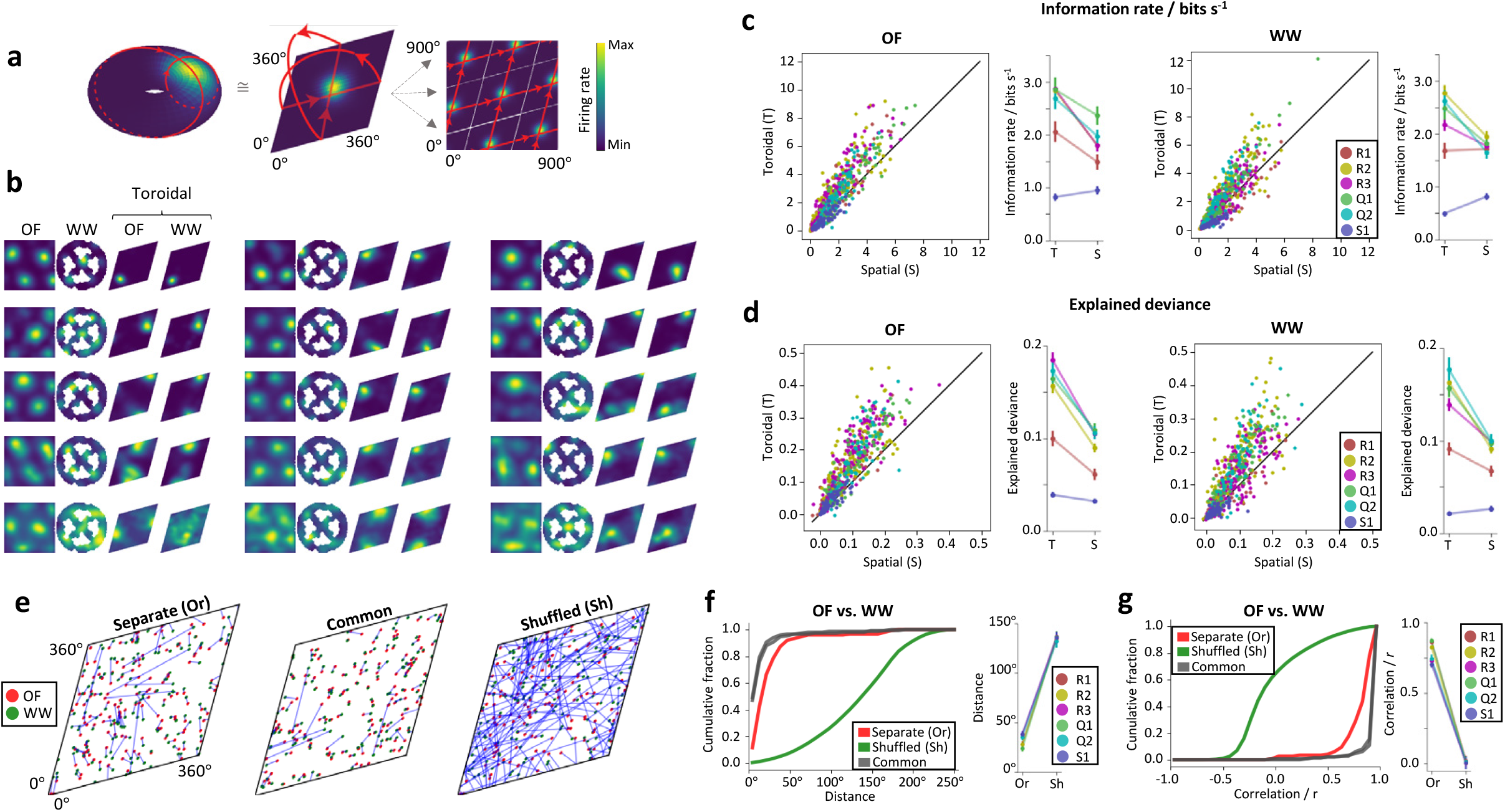
Cohomological decoding of position on an inferred state space torus. **a.** Individual grid cells have distinct firing fields on an inferred torus. Toroidal coordinates for population activity vectors may be decoded by computing time-varying, circular coordinates from the two significant 1D holes (red circles) identified by barcodes in the persistent cohomology analysis (Fig. 2c, d; Extended Data Fig. 5a). The Cartesian product of these circular coordinates localizes neural activity in the two-dimensional toroidal state space (Extended Data Fig. 4). Left: 3-D embedding of the toroidal state space displaying mean firing rate of a single grid cell as a function of toroidal position, colour-coded as indicated by scale bar. Middle: A 2-D torus may be formed by gluing opposite sides of a rhombus. Right: Arranging multiple rhombi to tesselate a 2-D surface reveals a grid-like pattern in the activity of the grid cell, akin to its spatial firing. **b.** Tuning of representative grid cells from module R2 to coordinates on the inferred torus. The panel shows every 10^th^ cell in a distribution where the cells are ranked according to their spatial information content (total of 15 cells are shown; see Extended Data Fig. 9 for the full set of data). Each row of 4 plots corresponds to one cell. From left to right: Rate maps showing firing rates across the spatial environment (open field, OF; wagon-wheel track, WW) and on the inferred torus (toroidal OF, toroidal WW). Firing rates are colour-coded as in **a**. The toroidal parametrizations were aligned to common axes before creating the rate maps. Note that firing locations on the torus are consistently preserved between OF and WW. **c.** Toroidal information rate and explained deviance in OF (left half) and WW (right half). Left of each: Scatterplots showing information rate for position on the inferred torus (T) vs position in the physical environment (S). The torus was inferred from WW for OF and from OF for WW; random noise was added to the T coordinates to compensate for tracking error in the S coordinates. Dots show individual cells; modules are colour-coded (inset). Right of each half: Mean information rate (±S.E.M) for position in S and T coordinates for each module. Note that 5/6 modules during OF and 4/6 during WW show higher information for the torus than for position. **d.** Scatterplots comparing the goodness-of-fit of two GLM models, based either on position in the spatial environment (S) or on the torus (T). We fitted a Poisson GLM model to the spike count using 3-fold cross validation. Goodness-of-fit is expressed as explained deviance, an *R*^2^-statistic whose values range from 0 to 1, with higher values indicating better fit. The two scatter plots show data from OF (left) and WW (right), respectively. Toroidal coordinates were processed as in **c.** Symbols as in **c**. Line diagrams to the right: Mean deviance explained for each module (±S.E.M.). In 6/6 modules from OF and 5/6 from WW, the explained deviance was larger for the toroidal regressor than the spatial regressor. **e.** Distribution of centres-of-mass for grid-cell firing fields on the torus (all cells of R2; OF, red; WW, green). Blue lines connect red and green peaks of the same cell (occasionally extending over the periodic boundaries of the toroidal sheet). Note the proximity of red-green pairs in the original data (after separate alignment of the two toroidal parametrizations “Separate” in left panel, or using the parametrization in OF for decoding in both OF and WW, “Common”, in middle panel) compared to shuffled data where rate maps are reordered randomly (right panel). **f.** Left: Cumulative frequency distribution showing distances between field centres on the torus for grid cells in module R2, in OF and WW (as in **e;** red: separate parametrizations “Separate”; grey: same parametrizations “Common”). Note that there are two grey curves (one parametrization for each environment). Values range between 0 and 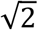 · 180° (the maximum possible distance between two points on the toroidal sheet). Green curves show distribution from 1,000 shuffled versions of the data. Right: mean distance between field centres (±S.E.M.) for each module (modules are colour-coded; Or, original data; Sh, shuffled data). Note substantially shorter distances for the original spike trains. **g.** Left: Cumulative frequency distribution showing Pearson correlations between the toroidal rate maps in OF and WW for grid cells in module R2 (colour-coding as in **f**). Right: Mean correlations (±S.E.M.) for each module; symbols as in **f**. Note substantially larger correlations for the original spike trains.

To test how faithfully location in the environment is mapped onto the toroidal representation, we asked whether grid-cell activity is predicted better by the cells’ tuning to the inferred torus than by their tuning to physical space. In each environment (OF or WW), we derived toroidal coordinates from the toroidal parametrization of the other environment (WW for OF sessions, OF for WW sessions), allowing us to dissociate the neural activity used to identify the torus from the neural activity to be described. In comparing tuning to toroidal and physical space, we added random gaussian noise to the toroidal position to compensate for errors in position tracking. For each cell, we compared the selectivity of its tuning, first to physical space, and second to the torus (with the cell in question omitted when decoding the toroidal coordinates). For 5/6 grid modules in OF and 4/6 in WW, the rate of information, in bits per second, that the cell’s firing rate conveyed about position was higher for position on the torus than in physical space (Fig 3c; R2, R3, Q1 and Q2: all *P* < 0.001 in OF and WW; R1: *P* < 0.001 in OF, *P* = 0.572 in WW; S1: *P* = 1.000 in OF and WW; Wilcoxon sign-rank test). We verified this difference by comparing the cross-validated prediction of two Poisson GLM-based encoding models of each cell’s activity that included toroidal position (decoded as above) and 2-D spatial position. For both environments (OF and WW), the toroidal covariate was closer to a perfectly fitted model of the data than the physical position covariate in 5/6 grid cell modules (Fig 3d; R1, R2, R3, Q1 and Q2: *P* < 0.001 for all OF and WW sessions; S1: *P* < 0.001 for OF, *P* = 1.000 for WW; Wilcoxon signed-rank test). Taken together, these differences point to toroidal structure as the primary feature of the population activity of grid cells, superior to that of the 2-D coordinates of the animal’s current position in the physical environment.

If grid cells operate on a toroidal manifold determined by intrinsic network features, this manifold may be expressed universally across environments, independently of sensory inputs. In this scenario, the toroidal selectivity for individual cells – a property specified by fixed connectivity in the CAN model – should also be preserved across environments. We tested this proposition by assessing, on the inferred tori, whether the locations of firing fields of different grid cells were maintained between OF and WW (Fig. 3b, Extended Data 9). To compare the toroidal parametrizations, we first aligned the axes of the toroidal coordinates before performing two sets of analyses. In one of them, we compared, for each cell, the distance between the centres of mass of the toroidal rate maps in OF and WW (Fig. 3e, f, Extended Data Fig. 5b, c). This distance was substantially shorter (mean ±S.E.M. of mean distances for all modules: 31.5±6.3 degrees) than that of control data in which the order of the rate maps in one environment was shuffled (135.8±1.7 degrees; max. possible distance 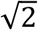 · 180 ≈ 254.6 degrees; data vs. shuffled: *P* < 0.001 in all modules; Wilcoxon signed-rank test). In the other analysis, we calculated the pairwise Pearson correlations of binned toroidal rate maps across the two environments (Fig. 3g; Extended Data Fig. 5d). Consistent with the centre-of-mass comparison, the correlations between OF and WW were higher in the observed data (mean±S.E.M. of mean *r*-values for all modules: 0.79±0.07) than the shuffled data (*r* = 0.01±0.01; *P* < 0.001 for all modules). Very similar results were obtained when applying the toroidal map from the same environment (either OF or WW) to activity from both environments, without applying any adjustment to the alignment of the toroidal axes (Fig. 3f, g, 16.0±3.4 degrees; *r* = 0.95±0.02; *P* < 0.001 for all modules and both mappings). Taken together, these findings suggest that single-cell tuning to toroidal coordinates is maintained across environments, and that physical space is mapped onto the same internal low-dimensional manifold irrespective of the specific environment.

### Persistence of toroidal topology during sleep

If population activity is mapped onto the same toroidal manifold independently of sensory inputs, the toroidal topology should be maintained also during sleep. To test this idea, the rats rested in a high-walled, opaque box placed in the centre of the OF. Periods of rapid-eye-movement (REM) sleep and slow-wave sleep (SWS) were identified based on the low-frequency rhythmic content of the aggregated multi-unit activity in combination with prolonged behavioural immobility (Extended Data Fig. 8; Online methods).

We started out with REM sleep. In 5 of the 6 grid modules, the persistent cohomology analyses of the point clouds of population activity identified topologies consistent with that of a torus, mirroring those of the OF and WW tasks (modules R1, R2, R3, Q1 and Q2; Fig. 4a, Extended Data Fig. 5a). In the barcode diagrams for these 5 modules, the lifetimes of one bar in 0-D, two bars in 1-D, and one bar in 2-D were substantially longer than the other bars of the point cloud, and exceeded those of a distribution of shuffled versions of the same data (*P* < 0.002 for all 5 modules). In the remaining module (S1), there were no long-lived bars in dimension 1 or 2 (Extended Data Fig. 5a.v), indicating an absence of toroidal structure during REM sleep. The lack of toroidal topology may partly reflect a low cell number (72 grid cells); approximately 60 simultaneously recorded cells were required to detect the torus with probability greater than 0.5 in downsampled data from module R2 (149 grid cells; Extended Data Fig. 3e). The barcode results were supported by the toroidal parametrization analyses, which revealed sharply tuned firing fields on the torus for these 5 grid modules (99.3±1.6% of the grid cells in each module had higher information rates than shuffled data, and in 95.3±7.2% of cells the toroidal tuning explained the activity better than a null model that assumes a constant firing rate; Fig. 4c). In addition, the toroidal tuning fields of the awake sessions maintained their relative locations in REM sleep (Fig. 4b, d, Extended Data Fig. 5b). The mean distance (±S.E.M.) between the peak locations of the pairs of rate maps from OF and REM (31.5±15.4 degrees) was far below the range of mean distances obtained in the shuffled data (Fig. 4e, Extended Data Fig. 5c; 135.8 ±2.3 degrees; *P* < 0.001 for all 5 modules), and mean pairwise correlations of toroidal rate maps for REM sleep and OF were substantially larger than in shuffled versions of the data (Fig. 4f, Extended Data Fig. 5d; *r* = 0.80±0.15 vs. 0.01±0.01; *P* < 0.001 for all 5 modules). We obtained similar results when comparing toroidal tuning of the grid cells in OF and REM to the same torus (from either REM or OF) (Fig. 4e, f; 19.5±10.9 degrees; *r* = 0.91+-0.07; *P* <0.001 for all 5 modules and both mappings). Taken together, these observations point to a preservation of toroidal structure from awake states to REM sleep.

**Figure 4:**
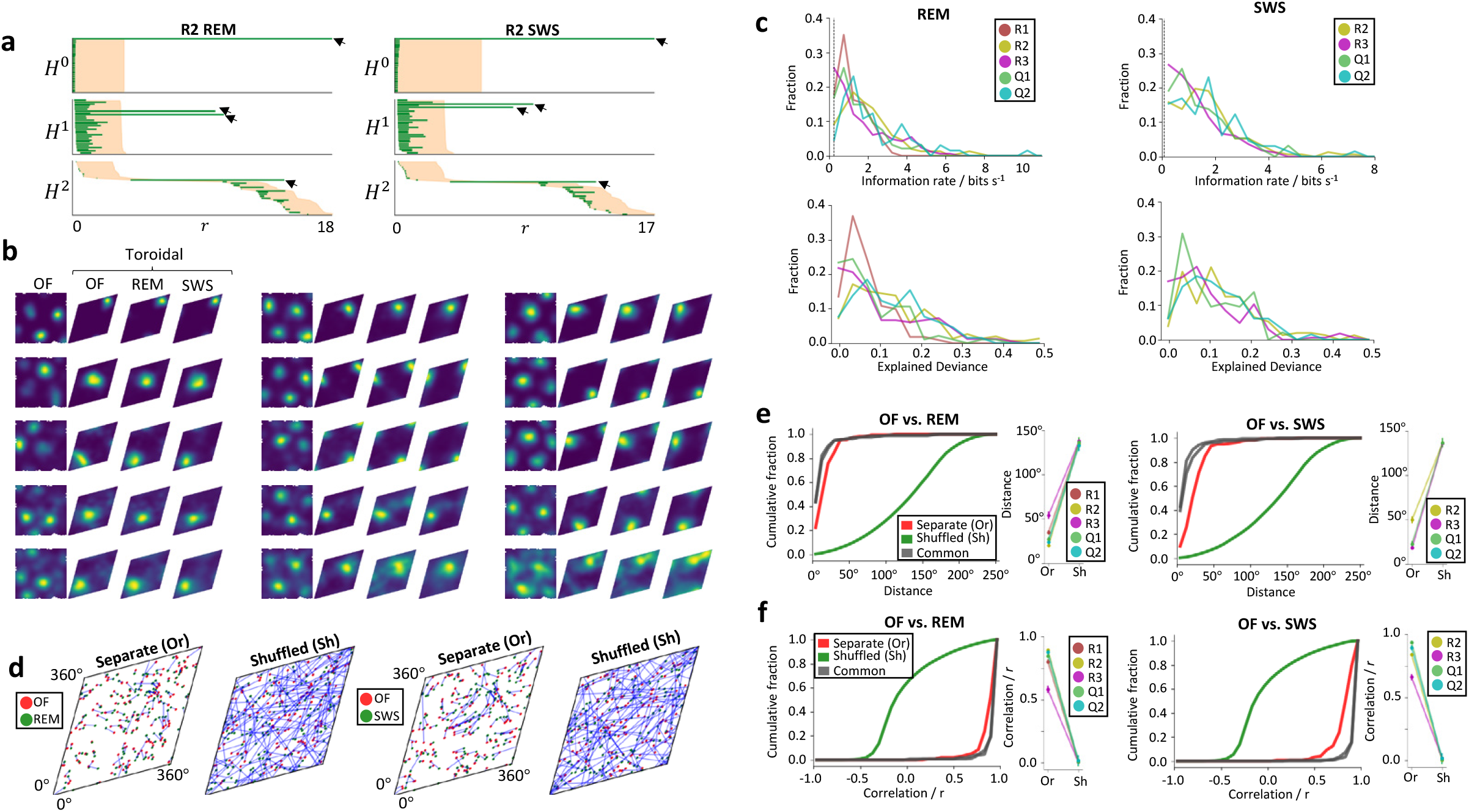
Preservation of toroidal structure during sleep when sensory inputs from the environment are minimized. **a.** Barcodes for grid-cell module R2 during REM sleep and SWS (as in Fig 2c, d for awake states). Arrows point to bars with lifetimes exceeding all values of the shuffled distribution: one bar in 0-D, two in 1-D and one in 2-D, as would be expected for toroidal topology. **b.** Toroidal rate maps showing preserved toroidal tuning relationships for individual cells across environments and brain states (showing every 10^th^ cell from a distribution ranked according to spatial information value of the cell’s toroidal rate map). From left: rate map for OF in physical coordinates, same OF session in toroidal coordinates, REM sleep in toroidal coordinates, and SWS in toroidal coordinates. Colour coding as in Fig. 3a. Toroidal alignment as in Fig. 3b. The full set of cells is shown in Extended Data Fig. 9. Tuning curves on the torus maintain relative locations between all conditions. **c.** Top: Histograms of the rate of information carried by individual cells’ activity about position on the inferred torus during REM (left) and SWS (right). Counts (fractions of the cell sample) are shown as a function of information rate (in bins of 0.5 bits/s) for all grid modules (colour-coded). The vertical dashed line (close to zero) shows mean information rate for shuffled distributions. The majority of cells have a higher information rate. Bottom: Explained deviance of a GLM model fitted to the spike count with the toroidal coordinates during REM (left) and SWS (right) as regressor. Distributions show counts (fractions of the cell sample) as a function of explained deviance, in bins of 0.035, for all grid modules. Values larger than 0 indicate that the fitted model explains the data better than a null model that assumes a constant firing rate. **d.** Distribution of field centres in toroidal rate maps for all grid cells in R2 during OF foraging (red) and sleep (green), as in Fig. 3e. Left pair: OF vs. REM; right pair: OF vs. SWS. Original data to the left in each pair, shuffled data to the right. **e.** Cumulative distributions showing distance between toroidal field centres for grid cells in module R2, as in Fig. 3f, but comparing awake behaviour in OF with REM (left) and SWS (right). To the right of each plot is shown the mean distance ±S.E.M. for all modules. The similarity between awake foraging and sleep indicates that toroidal topology is preserved. **f.** Cumulative distributions of Pearson correlation *r*-values of toroidal rate maps of grid cells in module R2, as in Fig. 3g, but comparing awake behaviour in OF with REM (left) and SWS (right). To the right of each plot is shown the mean *r*-value ±S.E.M. for all modules.

Toroidal topology was similarly maintained during SWS for 4 of the 6 grid modules (all except R1 and S1), with distinguishable bars corresponding to one 0-D bar, two 1-D bars and one 2-D bar (Fig. 4a and Extended Data Fig. 5a). Their lifetimes exceeded the persistence of any other bar in the barcode as well as the entire range of shuffled versions of the same data (P < 0.002 for each of the 4 modules). In agreement with the barcode analyses, strong tuning to toroidal coordinates was also expressed by individual cells, manifested in sharply tuned firing fields on the SWS torus (Fig. 4b, c, Extended Data Fig. 9; information rates showing 99.1±1.8% of the grid cells in each module were more informative of location on the torus than shuffled data; increase in explained deviance over the null model for 98.6±2.4% of cells). The spatial arrangement of toroidal firing locations in different cells during SWS matched that of awake and REM states (Fig. 4b, d, Extended Data Fig. 5b). The mean distance between peak firing locations during SWS and OF (29.8±14.3 degrees) was significantly shorter than in the shuffled data (Fig. 4e, Extended Data Fig. 5c; 135.8 ±2.3 degrees; *P* < 0.001 for all 4 modules), and pairwise correlations between SWS and OF toroidal rate maps were significantly higher (Fig. 4f, Extended Data Fig. 5c; *r* = 0.83±0.12 vs. *r* = 0.01 ±0.01; *P* < 0.001 for all 4 modules). Similar results were seen when using the toroidal parametrization in one of the conditions to decode the activity in both (Fig. 4e, f, Extended Data Fig. 5d; 17.8±9.2 degrees; *r* = 0.93+-0.06; *P* < 0.001 for all 4 modules and both mappings). Thus, the toroidal structure is maintained in both sleep conditions, despite the lack of external spatial inputs.

### Toroidal topology is confined to a major class of grid cells

Why was toroidal structure not visible during SWS in modules R1 and S1 (Fig. 5a and Extended Data Fig. 5a.v)? We asked whether this lack of toroidal structure reflected heterogeneity in the composition of the module. For module R1, an analysis of the activity correlation structure (Online methods) identified three classes: C_1-3_, consisting of 64, 30 and 17 grid cells, respectively. These classes were characterized by strong pairwise correlations within the class and weak correlations between classes (Fig. 5b). Confining the persistent cohomology analysis to the most populous class (C_1_), we again obtained a barcode indicative of toroidal structure (Fig. 5c). For all three classes, subsequent toroidal parametrization showed preservation of toroidal tuning locations in all brain states (mean distances between rate map centres: from 21 to 53 degrees in recorded data, from 79 to 176 degrees in shuffled data, *P* < 0.001 for all classes and all brain states; mean correlation between rate maps: from *r* = 0.66±0.05 to 0.85±0.03 in recorded data, from *r =* −0.25±0.03 to 0.38±0.04 in shuffled data, *P* < 0.001 for all classes and all brain states), suggesting that all cells participate in the toroidal structure, but with higher selectivity in C_1_ than C_2_ and C_3_ (Fig. 5d). For module S1, with only 72 cells, we were unable to identify toroidal structure in any class during SWS or REM. This module did express toroidal topology in the awake states, although with lower information rate and explained deviance in the toroidal tuning compared to the other modules (Fig. 3c, d; *P* = 0.0278; binomial test).

**Figure 5:**
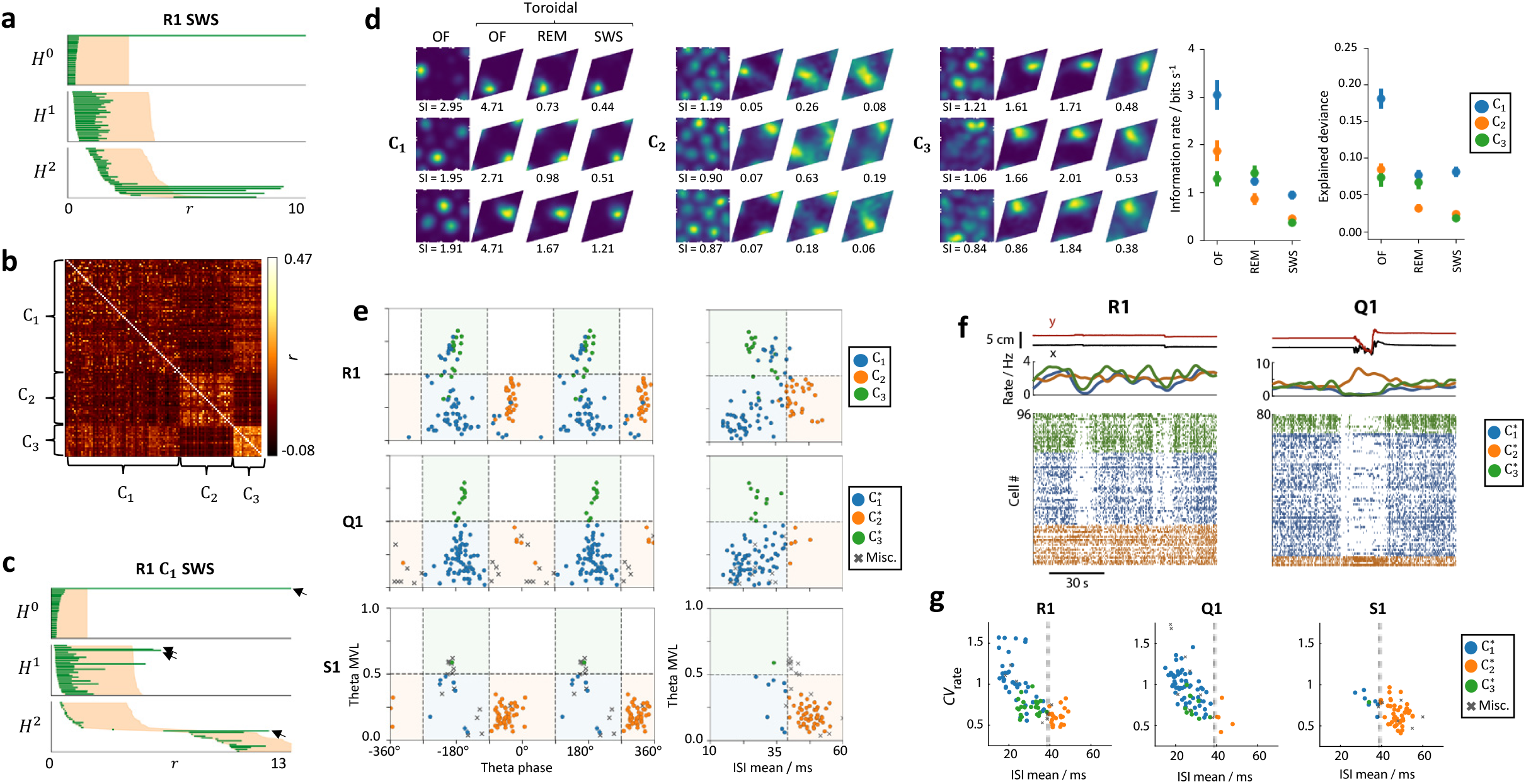
Subpopulations of grid cells with different temporal spiking statistics have different degrees of toroidal selectivity. **a.** The barcode of grid module R1, considered as a whole, does not show signs of toroidal structure during SWS. Symbols as in Fig. 2c. The barcode contains not one but several *H^2^* bars with lifetimes exceeding chance levels defined by shuffling. There is no *H^1^* bar with a significant lifetime. **b.** Correlation matrix showing pairwise correlation of firing rates for all grid cells belonging to R1. Correlation is colour-coded according to the scale bar. Complete-linkage clustering on the correlations reveals three clusters, C_1-3_, that each display strong inner correlation structure. Cluster boundaries are indicated on the axes of the correlation matrix. **c.** Barcode diagram indicating that C_1_ – the largest subpopulation – has toroidal structure. Arrows point to the most persistent features, as in Fig. 2c, d. The single long-lived *H°* bar, the two longest *H^1^* bars and the single prominent *H^2^* bar indicate toroidal structure. Note that a third *H^1^* bar exceeds the shuffled distribution, but its representation, given through cohomological decoding, did not show the striped pattern indicative of the circular axes of the toroidal structure (Extended Data Fig. 4). **d.** Decoding of toroidal coordinates for individual cells belonging to C_1-3_ (as in Fig. 4b), using toroidal coordinates from **c**. Left: Rate maps show the top three cells from a distribution ranked, for each class, on the cells’ spatial information (‘SI’) content. SI content is indicated beneath each spatial firing rate map; toroidal information rate is provided beneath each toroidal firing rate map). Toroidal coordinates are strictly preserved across states, as in Fig. 4b. The full set of cells is shown in Extended Data Fig. 9. Right: Mean (± S.E.M.) toroidal information rate and explained deviance scores for each class during OF, REM and SWS. Points are color-coded by class (C_1-3_). C_1_ shows higher mean toroidal selectivity than C_2_ (information rate: *P* < 0.05 REM and SWS; explained deviance: *P* < 0.05 all brain states; Kruskal-Wallis test, Dunn’s post-hoc test) and C_3_ (information rate: *P* < 0.05 OF and SWS; explained deviance: *P* < 0.05 OF and SWS). **e.** Grid cells segregate into classes with distinct spike timing characteristics. During OF foraging, the R1 correlation classes in **b** express differences in bursting probability and theta phase modulation. Upper Left: Scatter plot displaying, for individual cells (dots) in module R1, the relationship between cells’ assigned correlation classes (C_1-3_, colour-coded), their theta-phase modulation strength (mean vector length, MVL) and preferred theta firing phase. Theta-phase parameters were quantified in OF. Note the separation of classes: (1) C_1_ and C_3_ cells fire near the trough while C_2_ cells fire near the peak, and (2) C_3_ is more strongly modulated (higher MVL values) compared to C_1_ and C_2_. Upper Right: same as upper left, but with axes showing bursting probability during SWS (expressed as the mean interspike interval, “ISI mean”) and theta modulation strength in OF. Note the higher bursting probability (lower ISI mean) in C_1_ and C_3_ compared to C_2_. The shaded regions indicate the criteria subsequently used for classifying cells from all modules (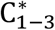; see also **f**, **g** and Extended Data Fig. 7). The two bottom rows show how these measures similarly separate cells of modules Q1 and S1, respectively. **f.** Burst-firing grid cells (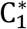 and 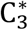) show coordinated low-activity states during rest (left, module R1; right, module Q1). The two plots show the activity of all grid cells in modules R1 (left) and Q1 (right) during a 100-second epoch of resting behaviour in the sleep box. Each row of dots shows the spikes of one grid cell, with cells grouped and coloured according to their assigned class. Above the raster plot are shown (upper) the animal’s 2-D position and (lower) the average smoothed firing rate of all cells in each class. Note the 10 – 30 s periods of rarefied activity visible in the raster plots which are only expressed by 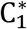 and 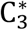 cells. Low-activity states sometimes coincide with brief arousals from sleep (as shown for Q1), but often occur within uninterrupted sleep (as shown for R1 here, and for S1 in Extended Data Fig. 7e). **g.** Bursting grid cells (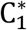 and 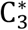) show more slow-timescale firing-rate variability during SWS. Each grid cell’s spike train was smoothed with a 2-second gaussian filter, and the coefficient of variation of the smoothed firing rate (*CV*_rate_) was calculated. Each scatter plot shows the distribution of grid-cell classes with respect to *CV*_rate_ and ISI mean (each dot represents one cell). The dot colours indicate the cells’ assigned class. The vertical grey line marks the mean ISI threshold used for classification in **e**. Note that *CV*_rate_ values are consistently lower for 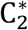 than for 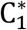 and 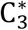. See Extended Data Fig. 7f for the remaining modules.

The confinement of toroidal topology to a distinct subset of grid cells in R1 led us to ask whether the R1 classes bore any relationship to previous accounts of heterogeneity in grid-cell firing characteristics^65–67^. We noted that the three grid-cell classes of R1 differed in theta phase-locking strength, preferred theta phase, and burst-firing probability, with less theta phase locking (but with preferential spiking at the trough) and more frequent bursting (shorter interspike intervals, ISIs) in C_1_ than C_2_ and C_3_ (Fig. 5e top row). To determine if similar structure was present in other data, we classified cells from all modules based on these measures (see Online methods), yielding three classes which we here refer to as 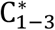, based on their correspondence to C_1-3_ in R1 (Fig. 5e, bottom two rows). Grid orientation and spacing of cells in the three classes were similar (Extended Data Fig. 7b), suggesting there was no difference in module identity between the classes. We found that modules with clear toroidal structure across behavioural conditions (R2, R3, Q1 and Q2) predominantly comprised 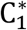 cells (76%, 67%, 71% and 82% of cells, respectively; Extended Data Fig. 7c). Conversely, in module S1, which failed to demonstrate toroidal topology in SWS, 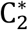 constituted 67% of cells. These cells resembled the subset of R1 cells that showed weaker toroidal tuning, weaker theta-phase modulation, with a preference for the theta peak, and low bursting probability (long ISIs). The preferential association of toroidal structure with 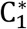 was also evident in the toroidal tuning scores of individual cells (information rate and explained deviance) in all brain states (OF, REM, SWS) in the five modules that expressed invariant toroidal topology (information rate: 15/15 module-by-experimental-condition combinations, *P* = 7 × 10^-8^, binomial test; explained deviance: 15/15 combinations, *P* = 7 × 10^-8^, binomial test; Extended Data Fig. 7d).

Finally, in each animal we observed brief low-activity population states^68,69^, during which the average firing rates of 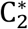 cells sometimes diverged strongly from 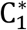 and 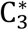 for periods of many seconds during rest. Particularly, bursting cells (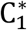 and 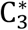) showed discrete periods of near-silence lasting 10–30 s, during which 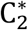 cells continued to fire at normal rates (Fig. 5f, Extended Data Fig. 7e). To quantify this phenomenon, we measured the variability of each grid cell’s firing rate on this slow timescale by calculating the coefficient of variation of the smoothed firing rate (*CV*_rate_) across the entire rest session (Online methods). In accordance with the persistence of 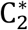 activity during low-activity periods, we found that the 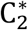 class showed the lowest median *CV*_rate_ among the three classes in all of the five modules which contained 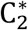 cells (Fig. 5g and Extended Data Fig. 7f; *P* = 0.00411, binomial test). Collectively, these observations suggest that there are distinct functional classes of grid cells, and that the toroidal neural code is preferentially maintained by a population of burst-firing, weakly theta-peak-preferring grid cells with high coefficients of variation during rest.

## Discussion

Our findings, from many hundreds of simultaneously recorded grid cells, show that the neural activity in grid cell populations invariably spans a manifold with toroidal topology, with movement on the torus matching the trajectory of the animal in the environment. The toroidal topology is not simply inherited from the encoded variable, since 2D space is not characterized by toroidal topology, as opposed, for example, to pitch and azimuth of head orientation, which together span a torus and thus naturally map in bats to a neural code with matching topology^70^. Using cohomological decoding to uncover the response properties of each cell on the toroidal manifold, we were able to demonstrate, in each environment and in both sleep and awake states, that the toroidal coordinates of the grid cells in an individual grid module were maintained, independently of external sensory inputs and possible environment-induced deformations of hexagonal symmetry in individual firing rate maps^4,61–64^. The uniform toroidal structure of the grid-cell manifold identified in the present work, across environments and behaviours, suggests that such distortions must occur in the mapping between physical space and the toroidal grid code rather than in the grid code itself.

The invariance of the toroidal manifold across environments and brain states is informative about the mechanisms that underlie grid cell activity. While toroidal topology can be generated by both CAN^1,13–16^ and feedforward^17–22^ mechanisms, the persistence of an invariant toroidal manifold across environments and behavioural states, under conditions that give rise to changes in the correlation structure of place-cell activity in the hippocampus^11,12^, is predicted only by CAN models. In these models, the topology is enforced by recurrent connectivity within the MEC, although external inputs may transiently and weakly perturb population activity patterns from the attractor. Previous work has shown that spike correlations persist across a variety of conditions in pairs of grid cells^9–12^, consistent with CAN models. However, pairwise correlations provide limited insight into the joint activity of grid cell populations, and it has remained unconfirmed whether this activity resides on an invariant, low dimensional manifold, and what the topology of such a manifold might be under circumstances where the regularity of the grid pattern is broken in individual cells. Here, using large-scale neural recordings combined with topological data analysis, we were able to put the hypothesis of a persistent toroidal manifold to test. The precise preservation of toroidal topology, and the maintained tuning relationships of individual grid cells to toroidal coordinates, provide strong support to the CAN hypothesis. While these findings do not exclude co-existing feedforward mechanisms, by which spatially periodic firing emerges under certain circumstances, e.g. early in development before the full maturation of recurrent connectivity^71–73^, the present observations point to intrinsic network connectivity as the mechanism underlying the rigid toroidal dynamics of the grid-cell system.

Finally, the findings suggest that toroidal topology is not always expressed uniformly across all grid cells of the module. In two of the six modules in our sample, a toroidal manifold was not identified during sleep when all grid cells were examined as a whole. One reason for this variation was the existence of multiple functional classes of grid cells within the grid module. These classes differed in burstiness, modulation by theta oscillations and variability of firing during sleep, with the strongest toroidal topology appearing in populations with high burstiness, a weak preference for the trough of the theta oscillation, and extended periods of near silence during sleep. Grid cells with these characteristics exhibited stronger toroidal tuning in all modules. Such heterogeneity in the basic firing properties of grid cells has been described previously^65–67^ but here we link these subtypes of grid cells to distinct population manifolds. The absence of clear toroidal topology in at least one class of grid cells (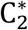 cells) may indicate a relative lack of CAN dynamics in this subpopulation, raising questions as to how this grid representation might be generated and maintained, and how it interacts with grid cells that exhibit stricter toroidal topology.

## Methods

### Subjects

The data were collected from 3 experimentally naive male Long Evans rats (Rats Q, R, and S, 300 – 500 g at time of implantation). The rats were group-housed with 3–8 of their male littermates prior to surgery and were singly housed in large Plexiglas cages (45 x 44 x 30 cm) thereafter. They were kept on a 12 hr light/12 hr dark schedule, with strict control of humidity and temperature. All procedures were performed in accordance with the Norwegian Animal Welfare Act and the European Convention for the Protection of Vertebrate Animals used for Experimental and Other Scientific Purposes.

### Electrode implantation and surgery

The rats were implanted with Neuropixels silicon probes^53,54^ targeting the MEC/PaS region. Two animals were implanted bilaterally with prototype Neuropixels ‘phase 3A’ single-shank probes and with one probe targeting MEC/PaS in each hemisphere; the third animal was implanted with a prototype Neuropixels 2.0 multi-shank probe in the left hemisphere. Probes were inserted at an angle of 25 degrees from posterior to anterior in the sagittal plane. Implantation coordinates were AP 0.05–0.3 mm anterior to the sinus and 4.2–4.7 mm lateral to the midline. The probes were inserted to a depth of 4200-6000 μm. The implant was secured with dental cement. The detailed implantation procedure has been described elsewhere^11,54^. After surgery, the animals were left to recover for approximately 3 hours before beginning recording. Postoperative analgesia (meloxicam and buprenorphine) was administered during the surgical recovery period.

### Recording Procedures

The details of the Neuropixels hardware system and the procedures for freely moving recordings have been described previously^53,54^. Briefly, electrophysiological signals were amplified with a gain of 500 (for phase 3A probes) or 80 (for 2.0 probes), low-pass-filtered at 300 Hz (phase 3A) or 0.5 Hz (2.0), high-pass-filtered at 10 kHz, and then digitized at 30 kHz (all steps performed by the probe’s on-board circuitry). The digitized signals were multiplexed by an implant-mounted “headstage” circuit board and were transmitted along a lightweight 5 m tether cable, made either using micro-coaxial (phase 3A) or twisted pair (2.0) wiring.

3-D motion capture (OptiTrack Flex 13 cameras and Motive recording software) was used to track the animal’s head position and orientation, by attaching a set of five retroreflective markers to implant during recordings. The 3-D marker positions were projected onto the horizontal plane in order to yield the animal’s 2-D position and head direction. An Arduino microcontroller was used to generate digital pulses which were sent to the Neuropixels acquisition system (via direct TTL input) and the OptiTrack system (via infra-red LEDs), in order to permit precise temporal alignment of the recorded data streams.

### Behavioural Procedures

Data were obtained from 4 recording sessions performed within the first 72 hours after recovery from surgery. The recordings were performed while animals engaged in three behavioural paradigms, each in a different arena within the same room. Abundant distal visual and sonic cues were available to the animal. On each day of recording, the animal remained continuously connected to the recording apparatus across the various behavioural sessions that were performed. Occasionally it was necessary to remove twists which had accumulated in the Neuropixels tether cable. In such cases, the ongoing behavioural task was paused while the experimenter gently turned the animal to remove the twists. During pre-surgical training, the rats were food-restricted, maintaining their weight at a minimum of 90% of their free-feeding body weight. Food was generally removed 12–18 hr before each training session. Food restriction was not used at the time of recording.

#### Open-field foraging task

Animals foraged for randomly scattered food crumbs (corn puffs) in a square open-field (OF) arena of size 1.5 × 1.5 m, with black flooring and enclosed by walls of height 50 cm. A large white cue card was affixed to one of the arena walls (height same as the wall; width 41 cm; horizontal placement at the middle of the wall). At the time of the surgery, each animal was highly familiar with the environment and task (10 – 20 training sessions lasting at least 20 minutes each).

#### Wagon-wheel track foraging task

The wagon-wheel (WW) track task was designed to function as a 1-D version of the 2-D open-field foraging task. The track’s geometry comprised an elevated circular track with two perpendicular cross-linking arms spanning the circle’s diameter. The track was 10 cm wide and was bounded on both sides by a 1-cm-high lip. Each section of the track was fitted with a reward point, placed halfway between the two nearest junctions, in the center of the section. Each reward point consisted of an elevated well which could be remotely filled with chocolate milk via attached tubing. To encourage foraging behaviour, a pseudorandom subset of the wells (between one and four of the eight wells) was filled at a given time, and the animal was allowed to explore the full maze freely and continuously. Wells were refilled as necessary when the animal consumed rewards. Each animal was trained to high performance on the foraging task prior to the surgery (collecting at least 30 rewards within a 30-minute session). Training to this level of performance took 5-10 half-hour sessions.

#### Natural sleep

For sleep sessions, the animal was placed in a black acrylic ‘sleep box’ with a 40 x 40 cm square base and 80-cm-high walls. The black coating of walls was transparent to infra-red, which allowed the 3-D motion capture to track the animal through the walls. The bottom of the sleep box was lined with towels, and the animal had free access to water. During recording sessions in the sleep box, the main room light was switched on and pink noise was played through the computer speakers to attenuate disturbing background sounds. Sleep sessions typically lasted 2-3 hours, but were aborted prematurely if the animal seemed highly alert and unlikely to sleep.

### Spike sorting and single-unit selection

Spike sorting was performed with KiloSort 2.5^54^. Briefly, the algorithm consists of three principal stages: (1) a raw-data alignment procedure which detects and corrects for shifts in the vertical position of the Neuropixels probe shank relative to the surrounding tissue; (2) an iterative template-matching procedure which uses low-rank, variable-amplitude waveform templates to extract and classify single-unit spikes; (3) a curation procedure which detects appropriate template merging and splitting operations based on spike train auto- and cross-correlograms. Some customizations were made to the standard KiloSort 2.5 method to improve its performance on recordings from the MEC/PaS region, where there is particularly high spatiotemporal overlap of spike waveforms due to the high density of cells. Therefore, the maximum number of spikes extracted per batch in step 1 above was increased, as was the number of template-matching iterations in step 2. To improve the separation between cells with very similar-looking waveforms, the upper limit on template similarity was raised from 0.9 to 0.975 in step 2 and to 1.0 on step 3, while supervising manually all merge and split operations from step 3, using a custom-made GUI running in MATLAB. The manual supervision ensured that Kilosort 2.5 did not automatically merge pairs of units with a dip in the cross-correlogram, which in our data was often due to out-of-phase spatial tuning. The merge and split operations were repeated several times to ensure the best separation between single units.

Single units were discarded if more than 1% of their interspike interval distribution consisted of intervals less than 2 ms. Additionally, units were excluded if they had a mean spike rate of less than 0.05 Hz or greater than 10 Hz (calculated across the full recording duration).

### Measurement of spatial position and direction tuning

During awake foraging sessions in the open-field arena or wagon-wheel track, only time epochs where the animal was moving at a speed above 2.5 cm s^-1^ were used for spatial or toroidal analyses. To generate 2-D rate maps for the open-field arena, position estimates were binned into a square grid of 3 × 3 cm bins. The spike rate in each position bin was calculated as the number of spikes recorded in the bin, divided by the time the animal spent in the bin. To interpolate the values of unvisited bins, two auxiliary matrices were used, *M_1_* and *M_2_*, setting visited bins equal to the value of the original rate map in *M_1_* and to 1 in *M_2_*, and setting unvisited bins to zero in both. One iteration of the image-processing ‘closing’ operation was then performed (binary dilation followed by erosion, filling out a subset of the non-visited bins) on *M_2_*, using a disk-shaped structuring element, first padding the matrix border by one bin. Both matrices were then spatially smoothed with a Gaussian kernel of smoothing width 2.75 bins. Finally, the rate map was obtained by dividing *M_1_* by *M_2_*. Rate-map spatial autocorrelograms and grid scores were calculated as described previously^55^. The selectivity of each cell’s position tuning was quantified by computing its spatial information content^74^, measured in bits per spike (see “Information rate and content”).

Head-direction tuning curves were calculated by binning the head direction estimates into 6° bins. The spike rate in each angular bin was calculated as the number of spikes recorded in the bin divided by the time the animal spent in the bin. The resultant tuning curve was smoothed with a Gaussian kernel with σ = 2 bins, with the ends of the tuning curve wrapped together. The selectivity of head-direction tuning was quantified using mean vector length (MVL) of the tuning curve. This was calculated according to:

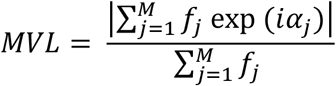

where vector *f* represents the tuning curve values (firing rates), vector α represents the corresponding angles, *M* is the number of tuning curve values, and |·| represents the absolute value of the enclosed term.

### Grid module classification

A novel method was implemented to detect populations of cells corresponding to grid modules by finding clusters of cells which expressed similar spatially periodic activity in the open field (Extended Data Fig. 2). Contrary to previous clustering-based methods for grid modules^5^, this approach makes no assumptions about the specific geometry of the grid pattern, thus making it less susceptible to the detrimental effects of geometric distortions such as ellipticity^5,61^.

For each MEC/PaS cell in a given recording, a coarse-resolution rate map of the open-field session was constructed, using a grid of 10 × 10 cm bins, with no smoothing across bins. The 2-D autocorrelogram of this rate map was calculated, and the central peak was removed by excluding all bins located < 30 cm from the autocorrelogram center. Bins located > 100 cm from the autocorrelogram center were also excluded. The autocorrelograms for all cells were subsequently converted into column vectors, *z*-standardized, then concatenated to form a matrix with spatial bins as rows and cells as columns. The nonlinear dimensionality reduction algorithm UMAP^75,76^ was then applied to this matrix, yielding a 2-dimensional point cloud where each data point represented the autocorrelogram of one cell (Extended Data Fig. 2a-d; UMAP hyperparameters: metric=‘manhattan’, ‘n_neighbors’=5, ‘min_dist’=0.05, ‘init’=‘spectral’). In the resultant 2-dimensional point cloud, cells with small absolute differences between their autocorrelogram values were located nearby one another. The point cloud was partitioned into clusters using the DBSCAN clustering algorithm (MATLAB function ‘dbscan’, minimum 30 points per cluster, eta = 0.6–1.0). In every recording, the largest cluster was mainly composed of cells which either lacked strong spatial selectivity or were spatially selective but without clear periodicity. All remaining clusters contained cells with high grid scores, and with similar grid spacing and orientation (Extended Data Fig. 2a-d); cluster membership was therefore used as the basis for grid module classification. In one recording (rat ‘R’ day 1), two clusters were identified which had similar average grid spacing and orientation (labelled as ‘R1a’ and ‘R1b’ in Extended Data Fig. 2a-d), suggesting that they represented the same grid module. R1b appeared to comprise cells with higher field-to-field firing rate variability, accompanied by more irregularities autocorrelograms. These two clusters were merged together in subsequent analysis (in which the resultant cluster is called ‘R1’).

A subset of the cells which were assigned to grid module clusters by the above procedure were tuned to both location and head direction (conjunctive grid × head direction cells). These cells, which were defined as having a head-direction tuning curve with mean vector length above 0.3, were discarded from further analysis.

### Classification of sleep states and spectral analysis

SWS and REM periods were identified based on a combination of behavioural and neural activity, following previously described approaches^11,77,78^. First, sleep periods were defined as periods of sustained immobility (> 120 s with locomotion speed < 1 cm s^-1^ and head angular speed < 6° s^-1^). Qualifying periods were then subclassified into SWS and REM based on the amplitude of delta- and theta-band rhythmic activity in the recorded MEC/PaS cells. Spike times for each cell were binned at a resolution of 10 ms and the resultant spike counts were binarized, such that ‘0’ indicated the absence of spikes and ‘1’ indicated one or more spikes. The binarized spike counts were then summed across all cells (Extended Data Fig. 8a). The rhythmicity of this aggregated firing rate with respect to delta (1-4 Hz) and theta (5-10 Hz) frequency bands was quantified by applying a zero-phase, fourth-order Butterworth band-pass filter, then calculating the amplitude from the absolute value of the Hilbert transform of the filtered signal, which was smoothed using a Gaussian kernel with σ = 5 s and then standardized (“*z*-scored”). The ratio of the amplitudes of theta and delta activity was hence calculated (theta/delta ratio, ‘TDR’). Periods where TDR remained above 5.0 for at least 20 s were classified as REM; periods where TDR remained below 2.0 for at least 20 s were classified as SWS (Extended Data Fig. 8b).

Spectral analysis was performed on 10 ms-binned multi-unit activity using the multi-tapered Fourier transform, implemented by the Chronux toolbox (http://www.chronux.org, version 2.10, function “mtspectrumsegc”). Non-overlapping 5-second windows were used, with a frequency bandwidth of 0.5 Hz and the maximum number of tapers.

### Visualization of toroidal manifold

For each module of grid cells, spike times of co-recorded cells in the OF were binned for each cell at a resolution of 10 ms, and the binned spike counts were convolved with a Gaussian filter with σ = 50 ms. Time bins where the animal’s speed was below 2.5 cm s^-1^ were then discarded. To account for variability of average firing rates across cells, the smoothed firing rate of each cell was *z*-scored. For computational reasons, the time bins were downsampled, taking every 25^th^ time bin (equating to 250 ms intervals between selected samples). Collectively, the downsampled firing rates of the full population of cells formed a matrix with time bins in rows and cells in columns. PCA was applied to this matrix (treating time bins as observations and cells as variables), and the first six principal components were retained (Extended Data Fig. 3b,c). UMAP^75,79^ was then run on these six principal components (with time bins as observations and principal components as variables). The hyperparameters for UMAP were: ‘n_dims’=3, ‘metric’=‘cosine’, ‘n_neighbours’=5000, ‘min_dist’, 0.8 and ‘init’=‘spectral’.

For visualizing the toroidal manifold during WW, smoothed firing rates were first calculated by the same procedure described above for OF. Subsequently, to allow comparison of the toroidal manifold between OF and WW, the same PCA and UMAP transformations calculated for the OF data were re-applied to the WW data.

### Preprocessing of population activity for topological analyses

Each topological analysis was based on the activity of a single module of grid cells, during a single experimental condition in one recording session. The experimental conditions were: open field foraging (OF), wagon-wheel track foraging (WW), slow-wave sleep (SWS), and rapid eye-movement sleep (REM). Sleep epochs of the same type were collected from across the recording and concatenated for analysis purposes. Similarly, in one case (rat S), two WW task sessions were concatenated in order to increase the sample size.

In total there were 27 combinations of module (Q1, Q2, R1, R2, R3, S1) and experimental condition (OF day 1, OF day 2, WW, REM, SWS).

Preprocessing of spike trains began by computing delta functions centred on the spike times (valued 1 at time of firing, 0 otherwise), and convolving these temporally with a Gaussian kernel with σ = 50 ms (OF, WW and REM) or 25 ms (SWS). Samples of the smoothed firing rates of all cells (“population activity vectors”) were then computed at 50 ms intervals. The awake states were further refined by excluding vectors which originated from time periods when the animal’s speed was below 2.5 cm s^-1^.

Computing the persistent cohomology of a point cloud is computationally expensive and may be sensitive to outliers (e.g. spurious points breaking the topology of the majority of points in the point cloud). For this reason, it is common to preprocess the data by downsampling and dimension-reducing the point cloud. The same preprocessing procedure was used for all datasets in the present study.

First, the data points were downsampled by keeping the 15,000 most active population activity vectors (as measured by the mean population firing rate). The reduced point cloud was then *z*-scored and was projected to its 6 first principal components – thus reducing noise, while keeping much of the variance and grid structure (see Extended Data Fig. 3b-d). To further simplify the low-dimensional point cloud, a novel downsampling technique was introduced, based on a point-cloud density strategy motivated by a topological denoising technique introduced by Kloke and Carlsson (2009)^80^ and a fuzzy topological representation used in UMAP^75,81^. Parts of the open-source implementation of the latter were copied in this computation. This approach consisted of assigning, for each point, a neighborhood strength to its *k* nearest neighbors, and subsequently sampling points that represent the most tight-knit neighborhoods of the point cloud in an iterative manner. First, we defined 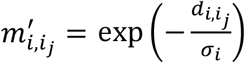 where 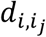 is the cosine distance between point *x*, and its *j-th* nearest neighbor and σ*_i_* is chosen to make 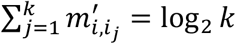, using *k* = 1500. The neighborhood strength was then obtained by symmetrizing: 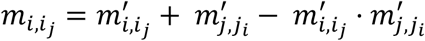. Finally, the point cloud was reduced to 1200 points by iteratively drawing the *i*-th point as: 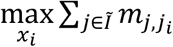, where *Ĩ* denotes the indices of the points not already sampled. In other words, for each iteration, the sampled point is the one with the strongest average membership of the neighborhoods of the remaining points.

To compute the persistent cohomology of the downsampled point cloud, the neighborhood strengths were first computed for the reduced point cloud (using *k* = 800) and its negative logarithm was taken, obtaining a distance matrix. This matrix was then given as input to the Ripser implementation^82,83^ of persistent cohomology, returning a barcode. In short, the barcode gave an estimate of the topology of the fuzzy topological representation of the six principal components of the grid-cell population activity. Thus, in essence, the first step of UMAP^75^ was applied before describing the resulting representation with persistent cohomology, instead of using it to project each point of the point cloud to a representation of user-specified dimensionality for visualization (Extended Data Fig. 3a5, a6). This gives a more direct and stable quantification of the global data structure, without having to choose an initialization^57^ or optimize a lower-dimensional representation.

### Persistent cohomology

Persistent cohomology is a tool in topological data analysis used to characterize the manifold assumed to underlie the data. It has previously been successful in analysing neural data, describing the ring topology of head direction cell activity^50–52^, the spherical representation of population activity in primary visual cortex^84^, and the activity of place cells^85–88^.

The general outline of the algorithm is: Each point in the cloud is replaced by a ball of infinitesimal radius, and the balls are gradually expanded in unison. Taking the union of balls at a given radius results in a space with holes of different dimensions. The range of radii for which each hole is detected is tracked; this is referred to as the “lifetime” of the hole and is represented by the length of a bar. The totality of bars is referred to as the barcode.

The software package Ripser^82,83^ was used for all computations of persistent cohomology. Ripser computes the persistent cohomology of ‘Vietoris-Rips complexes’ (which approximate the union of balls for different radii), constructed based on the input distance matrix and a choice of coefficients (in our case, ℤ_47_-coefficients), and outputs the barcode and cocycle representatives for all bars.

To verify that the lifetimes of prominent bars in the barcodes were beyond chance, shuffled distributions were generated for the persistence lifetimes in each dimension. In each shuffling, the spike train of each cell was shifted independently in time by rolling the firing rate arrays a random length between 0 and the length of the session. The same pre-processing and persistence analysis was then performed on the shifted spike trains as for the unshuffled data. This was performed 1,000 times, and each time a barcode was obtained. The barcodes were concatenated for all shuffles and the maximum lifetime was found for each dimension. This lifetime served as a significance criterion for the bar lifetimes. It is noted, however, that this is a heuristic and that statistics of barcodes is still not well established.

### Cohomological decoding

To further investigate the results identified by the barcode, the ‘cohomological decoding’ procedure introduced by De Silva et al (2011)^58^ was used to calculate a toroidal parametrization of the point clouds of population activity, assigning corresponding positions on each of the two circular features identified by the 1-D bars with the longest lifetime. This construction is motivated by the isomorphism between the first cohomology group of a topological space *X* with coefficients in Z and the set of homotopy equivalent maps from *X* to the circle (S^1^)^89^:

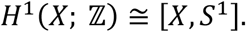

In the present case, persistent cohomology was first applied to the grid-cell population activity and took *X* to be the Vietoris-Rips complex for which the two longest-lived one-dimensional bars in the barcode (representing each of the two circles of the torus) existed. To define the desired toroidal coordinates on a domain that was as large as possible, we chose the complex given at the scale of the birth plus 0.99 times the lifetime of the second longest-lived one-dimensional bar in the barcode^50,58,90^. Next, the cocycle representatives (given by the persistent cohomology implementation of Ripser^82,83^), of each of the chosen 1-D bars, defined ℤ_47_-values for each of the edges in the complex. These edge values were then lifted to integer coefficients and subsequently smoothed by minimizing the sum over all edges (using the *scipy* implementation *lsmr*). The values on the vertices (points) of each edge followed from the edge values and gave the circular parametrizations of the point cloud. The product of the two parametrizations thus provided one map from the neural activity to the two-dimensional torus – i.e. giving a toroidal coordinatization (decoding) of the data.

Since persistent cohomology was computed for a reduced data set of 1200 points and therefore circular parametrizations were obtained only for this point cloud, each parametrization was interpolated to the population activity from the rest of the session(s). First, the 1200 toroidal coordinates were weighted by the normalized (“*z*-scored”) firing rates of the cells at those time points, obtaining a distribution of the coordinates for each grid cell. The decoded toroidal coordinates were then computed by finding the mass centre of the summed distributions, weighted by the population activity vector to be decoded. These activity vectors were calculated by first applying a Gaussian smoothing kernel of 15 ms standard deviation to delta functions centred on spike times, sampling at 10 ms intervals and then *z*-scoring the activity of each cell independently. Time intervals containing no spikes from any cell were subsequently excluded. When decoding was used to assess or compare the tuning properties of single cells (e.g. comparison of toroidal versus spatial description), the coordinates were computed using the weighted sum of the distributions of the other cells, i.e. the contribution of the cell to be assessed or compared was removed. When comparing preservation of toroidal tuning across two sessions, coordinates were interpolated using either the toroidal parametrization in each session independently (“Separate”) or using the same toroidal parametrization in both sessions (“Common”).

### Toroidal rate map visualization

For visualization, toroidal firing rate maps were calculated in the same way as the physical space covariate (see “Measurement of spatial position and direction tuning”), first binning the toroidal surface into a square grid of 7.2° × 7.2° bins and computing the average spike rate in each position bin. However, for toroidal maps, it was necessary to address the 60° angle between the toroidal axes before smoothing. After binning the toroidal coordinates, the rate map was “straightened” by shifting the bins along the *x*-axis (“horizontally”) the length of 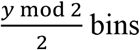, where *y* is the vertical enumeration of the given bin. Copies of the rate map were then tiled in a three-by-three square (similar to Fig. 3a right), before applying the closing and smoothing operations as for the spatial firing rate map. The single toroidal rate map was finally recovered by cutting out the center tile, rotating it 90 degrees and defining 15° shear angles along both the *x-* and *y*-axis to correct for the 60° offset between them.

### Comparison of spatial periodicity between environments

Differences in grid periodicity between OF and WW environments were quantified for a given cell by comparing the grid scores in the two behavioural conditions. Two alternative methods were used to generate the spatial autocorrelograms for this comparison: (1) comparing the autocorrelograms for OF and WW directly, and (2) comparing autocorrelograms for OF and WW after first equalizing the spatial coverage between the two conditions.

For method (1), rate maps were calculated as specified in the above section “Measurement of spatial position and direction tuning”, using the same grid of 3 × 3 cm bins for both environments. This set of bins spanned the entirety of the OF arena and covered most of the WW track apart from some small regions at the outer extrema, which were discarded for the purpose of this analysis. For each of the two rate maps, the autocorrelogram was computed and grid score calculated.

Method (2) was similar to method (1), except that the cell’s OF rate map was converted into a “masked OF” rate map, by removing all bins which were unvisited by the animal in the WW session. This effectively equalized the position coverage between the two conditions, and thus allowed for a more valid comparison.

### Comparison of toroidal versus spatial description

The explanatory significance of the toroidal description was evaluated by comparing statistical measures of how well the toroidal coordinates explained neural activity on the torus and in physical space. For a fair comparison, it was important to avoid overfitting, which might occur if a toroidal parametrization of a point cloud is used to describe that same set of data points. Two precautions were taken to avoid such overfitting: first, the data was decoded using the toroidal parametrization from a different condition (an OF session for a WW recording and a WW session for an OF recording), and second, the cell for which the statistical measurement was made was omitted from the decoding.

The comparison of toroidal and environmental representations also accounted for tracking error in the physical position estimate, which mainly resulted from the approximately 4 cm vertical offset of the tracking device above the animal’s head. This causes a discrepancy when the angle *α* between the animal’s zenith and the axis of gravitation is different from 0°, measured as 4 tan *α* cm. The mean discrepancy in the recorded position data was measured to 1.5cm. To account for this error of the position estimate, proportional Gaussian noise was added to the toroidal coordinates, using a standard deviation of 1.5cm/Ω, where Ω denotes the grid spacing of the particular grid-cell module, estimated from the mean period of the fitted cosine waves of the toroidal coordinates in the open field (see “Rhomboidal fitting and toroidal alignment”).

### Information rate and content

The information rate (*I_R_*) was calculated as previously described^74^, to quantify and compare the amount of information carried by single-cell activity about the location on the torus and physical space per second. Both covariates were binned in a *M* = 15×15 grid of square bins. For each bin *j*, the average firing rate *f_j_* (given in spikes per second), and the occupancy ratio, *P_j_*, were computed. The information rate for each grid cell was then given as:

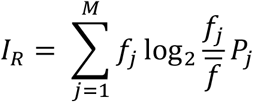

where 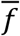 is the mean firing rate of the cell across the entire session. The information content per spike, *I_c_* was calculated by dividing the information rate by the mean firing rate:

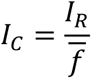

### Deviance explained

Deviance explained was computed to measure how well a Poisson GLM model fitted to the spike count was at representing the data, using either the toroidal coordinates or the tracked position as regressors. A similar setup was used to that of a previous study^91^, with a smoothness prior for the GLM to avoid overfitting.

Both the toroidal and spatial coordinates were binned into a 15 × 15 grid of bins, and GLM design matrices were built with entries *X_i_*(*t*) = 1 if the covariate at time *t* fell in the *i*-th bin and *X_i_* (*t*) = 0 otherwise.

The Poisson probability of recording *k* spikes in time bin *t* is:

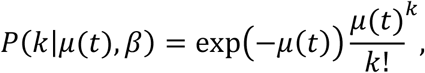

where 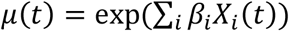 is the expected firing rate in time bin *t*. The parameters *β* of the Poisson GLM were optimized for each covariate by minimizing the cost function:

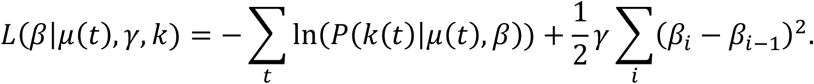

The first term is the negative log-likelihood of the spike count in the given time bin, while the second term puts a penalty on large differences in neighbouring parameters, enforcing smoothness in the covariate response of the predicted spike count.

*βs* was initialized to zero and performed the minimization of the loss function by first running two iterations of gradient descent, before optimizing using the *‘ l-bfgs-b’-algorithm* (as implemented in the *scipy.optimize-module*) with ‘gtol’=1e-5 as the cut-off threshold, and finally running two more iterations of gradient descent. A 3-fold cross validation procedure was used, repeatedly fitting the model to two thirds of the data and testing on the held-out last third.

The smoothness hyperparameter *γ* was optimized *a priori* on each grid cell module based on the summed likelihood, testing 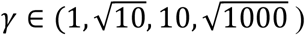, and found to be either 1 or 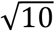 in all cases.

Similarly, after fitting a null model (using only the intercept term) and the saturated model (perfectly fitting each spike count), the deviance explained could be computed as:

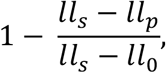

where *ll*_p_, *ll*_0_ and *ll_s_* denotes the cross-validated log likelihood of the fitted model, the null model and the saturated model, respectively. This provides a normalized comparison describing the difference between the fitted model and the idealized model.

### Rhomboidal fitting and toroidal alignment

The cohomological decoding of the torus gave two circular coordinates of arbitrary origin and orientation (i.e. clockwise or counterclockwise evolution). To infer a geometric interpretation of the tori and compare the toroidal parametrizations across modules and conditions, two cosine waves cos(*ωt* + *φ*) were fitted to the open-field mappings of the decoded circular coordinates. *t* is the center 100^2^-bins of a 540° × 540°-valued 150^2^-bin grid rotated *θ* degrees. The parameters (*ω, φ, θ*) were optimized by minimizing the square difference between the cosine wave and the cosine of the circular mean of the circular coordinates in 100^2^ bins of the physical environment (smoothed using a Gaussian kernel with 1 bin standard deviation). Estimates were first obtained by finding the minimum when testing all combinations of the evenly spaced values: ω ∈ [1,6], *φ* ∈ [0,360) and *θ* ∈ [0,180), using 10 steps for each interval. The parameters of the cosine wave were further optimized using the *slsqp*minimization algorithm (as implemented in the *scipy.optimize-* module using default hyperparameters). The period was computed as 1.5*m*/*ω*.

Next, to obtain the angle between the directions of the circular coordinates, the coordinate for which cosine of cos(*θ*) was largest was defined as the first coordinate (*θ*_1_), and the other was defined as the second coordinate (*θ*_2_). Although these two coordinates fully describe the toroidal location, the hexagonal torus allows for three axes, with the second and third axes respectively oriented at 60 and 120 degrees relative to the first, thus making each axis a linear combination of the other two. The difference in directions was given by *θ_1_* — *θ_2_*. However, to obtain similar axes of the different tori, the clockwise orientation of each circular coordinate was first determined by noting whether (*ωt* + *φ*) or 360°-(ωt + *φ*) best fit the spatial mapping of the circular means of the toroidal coordinates, and reoriented to obtain the same direction for both coordinates. Second, if the angle was greater than 90 degrees, the second coordinate was replaced with *θ_3_* = *θ_2_* + 60° · *θ*_1_. Finally, the origin of the coordinates was fixed by shifting each set around the circle by the mean angular difference between the given set of coordinates and the coordinates found during the open field session for each module.

For visualization of the rhombi (Extended data fig. 4), the angle of the vectors found by the cosine fit was similarly updated and furthermore it was necessary, in some cases, to rotate both vectors 30 degrees depending on whether one of the axes was directed outside of the box.

### Preservation of toroidal tuning

Centre-to-centre distance and Pearson correlation was computed between toroidal tuning maps of different sessions to measure the degree of preservation between the toroidal descriptions.

First, the preferred toroidal firing location for each cell was computed as the centre of mass of the toroidal firing distribution:

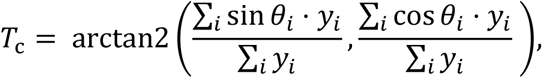

where *y_i_* denotes the mean spike count of the given cell in the *i*-th bin whose binned toroidal coordinates are given by *θ*_i_. The distance between mass centres found in two sessions (“5_1_” and “S_2_”) was then defined as:

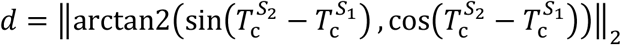

where ‖ · ‖_2_ refers to the L_2_-norm.

Pearson correlation between two tuning maps was computed by flattening the smoothed 2-D rate maps to 1-D arrays and calculating the correlation coefficient, r, using the *pearsonr-* function given in the *scipy.stats-library*.

To determine how much the preservation of the toroidal representations across two sessions (measured with Pearson correlation and peak distance) differed from a random distribution, the indices of the cells in one of the sessions before computing correlation and distance for the pair of conditions. This process was repeated 1,000 times, and the p-value was calculated from the rank of the original *r*-value or distance with respect to the shuffled distribution.

### Correlation clustering-based classification of grid cells within modules

The correlation coefficient matrix of the smoothed firing rates of all R1 cells was computed (Fig. 5b) and complete linkage clustering was applied on the correlations using the *scikit-learn* implementation *AgglomerativeClustering* with the *L*_1_-distance as affinity measure and ‘distance_threshold’ = 11.

### Theta phase tuning

To measure the modulation of a cell’s firing rate by the phase of theta oscillations, the method described in “Classification of sleep states and spectral analysis” was first used to calculate the 5–10 Hz bandpass-filtered multi-unit spiking activity. The instantaneous theta phase was then estimated by computing the angle of the complex-valued Hilbert transform of the filtered multi-unit activity. This method defined phases 0° and 180° respectively as the (approximate) maximum and minimum of neural activity. To simplify comparison with previous studies of MEC LFP theta waves, which are approximately in counterphase to levels of spiking activity in superficial MEC layers^92^, these phases were shifted by 180° to give a final definition of phase in which 0° and 180° respectively correspond to the conventional LFP “peak” and “trough” phases. Finally, the cell’s tuning curve with respect to theta phase was computed, and expressed as a mean vector length (MVL) score, according to the same method described for head-direction tuning curves (see “Measurement of spatial position and direction tuning”).

### Burst-firing probability

A cell’s probability of bursting was determined by measuring its mean inter-spike interval (ISI). ISIs were computed as the temporal difference of consecutive spikes. ISIs larger than 100 ms were excluded before calculating the mean.

### Classification of grid cells by theta-phase tuning and burst-firing probability

Grid cells from all modules were classified based on their theta-phase tuning (preferred phase and mean vector length, MVL) and burst-firing probability, calculated as described above. Classification criteria were chosen that provided good separation of the three classes of cells in R1 defined by the correlation clustering (Fig. 5b, e).

The classification thresholds were as follows: preferred theta phase *θ_a_* = —90° and *θ_b_* = 90°, theta MVL threshold *τ* = 0.5, and burst-firing probability *β* = 39ms, giving classes 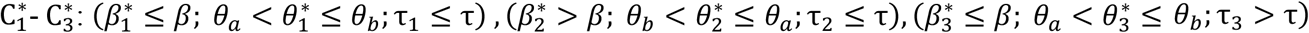. Cells that did meet the criteria for any class are referred to as ‘Miscellaneous’.

### Firing rate coefficient of variation

The coefficient of variation of the firing rate (*CV*_rate_) of a given cell was used as a metric for quantifying the slow fluctuations of the cell’s firing rate associated with suppression of activity during low-activity brain states^68,69^. *CV*_rate_ was computed by first binning the cell’s spike times at a resolution of 100 ms, then temporally smoothing the binned spike counts using a Gaussian window with σ = 2 s. All time bins which were not within SWS periods were discarded at this point, and hence the coefficient of variation was calculated from the remaining firing rate values according to the formula 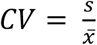, where s is the standard deviation of the firing rate, and 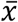 is the mean firing rate. Because estimates of *CV*_rate_ are noisy for small values of 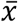, to increase the reliability of this analysis only cells with mean firing rates above 1 Hz were included.

### Minimum number of cells displaying toroidal structure

To address the question of how many cells are minimally needed to expect to see toroidal structure, random samples of *n* = 10, 20, 30 …, 140 cells were taken from R2 (*N* = 149 cells) during open field foraging, and the same topological analysis was repeated as for the whole population. The cells were resampled 1000 times for each number of cells in the subsample. To determine whether toroidal structure was detected, a heuristic was introduced based on the circular parameterization given by the two most persistent 1-D bars in the barcode mapped onto physical space. An estimate of the resulting planar representation of the torus was obtained by fitting planar cosine waves to each mapping (see “Rhomboidal fitting and toroidal alignment”). For the analysis to be determined ‘successful’ in detecting toroidal structure, we required: (i) the mean value of the least-squares fitting (across bins of the mapping) to be less than 0.25; (ii) the angle of the rhombus to be close to 60 degrees (between 50 and 70 degrees) and (iii) the side lengths to be within 25% of each other,.

### Simulated CAN Models

To confirm the expected outcomes of topological analyses of grid cell CAN models, grid cells were simulated using two different, noise-less CAN models (Extended Data Fig. 6).

First, a 56 × 44 grid cell network was simulated based on the CAN model first proposed in Burak & Fiete, 2009^13^, but using solely lateral inhibition (for details see Couey, 2013^15^) in the connectivity matrix, *W*. The animal movement was given as the first 1000 s of the recorded trajectory of rat ‘R’ during open field session, originally sampled at 10 ms, and interpolated to 2-ms time steps. The speed, *ν*(*t*), and head direction *θ*(*t*) of the animal was calculated as the (unsmoothed) displacement in position for every time step. The activity, *S*, was updated as:

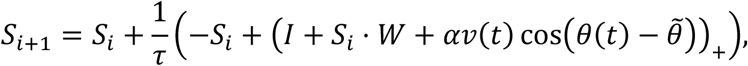

where (...)_+_ is the Heaviside function and *θ* is the population vector of preferred head directions. The following parameters were used: *i* = 1, *‘α’* = 0.15, *‘I’* = 2, *‘W_0_’* = −0.01, *‘R’* = 20 and *‘τ’* = 10, and let the activity pattern stabilize by first initializing to random and performing 2000 updates, disregarding animal movement. For computational reasons, the activity was set to 0 if *S_i_*, < 0.0001. The simulation was subsequently downsampled keeping only every 5^th^ time frame.

Next, a 20 × 20 grid cell network was simulated, for a synthetically generated open field trajectory (‘random walk’), based on the twisted torus model formulated by Guanella et al^16^. The parameter values and the code for computing both the grid cell network (choosing a single grid scale by defining the parameter ‘grid_gain’ = 0.04) and the random navigation (using 5000 time steps) were given by the implementation by Santos Pata^93^.

### Idealized Torus Models

To compare the results of both the original and simulated grid cell networks with point clouds where the topology is known, *a priori*, to be toroidal, points were sampled from a square and a hexagonal torus. First, a 50 × 50 (angle) mesh grid (*θ*_1_, *θ*_2_) was created in the square [0,2π) × [0,2π) and slight Gaussian noise (*ϵ* = 0.1 · *N*(0,1)) was added to each angle. The square torus was then constructed via the 4-D Clifford torus parametrization: (cos(*θ*_1_), sin(*θ*_1_), cos(*θ*_2_), sin(*θ*_2_). The hexagonal torus was constructed using the 6-D embedding: (cos(*θ*_1_), sin(*θ*_1_), cos(*a*_1_ *θ*_1_ + *θ*_2_), sin(*a*_1_ *θ*_1_ + *θ*_2_), cos(*a*_2_ *θ*_1_ + *θ*_2_), sin(*θ*_2_ *θ*_1_ + *θ*_2_)), where 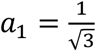 and 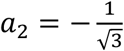.

## Data analysis and statistics

Data analyses were performed with custom-written scripts in Python and MATLAB. Samples included all available cells that matched the classification criteria for the relevant cell type. The study did not involve any experimental subject groups; therefore, random allocation and experimenter blinding did not apply and were not performed.

The most intensive computations were performed on resources provided by the NTNU IDUN/EPIC computing cluster^94^.

## Code availability

Code for reproducing the analyses in this article will be available after publication at Figshare and/or GitHub.

## Data availability

The datasets generated during the current study will be available after publication at Figshare.

---

**Extended Data Figure 1:**
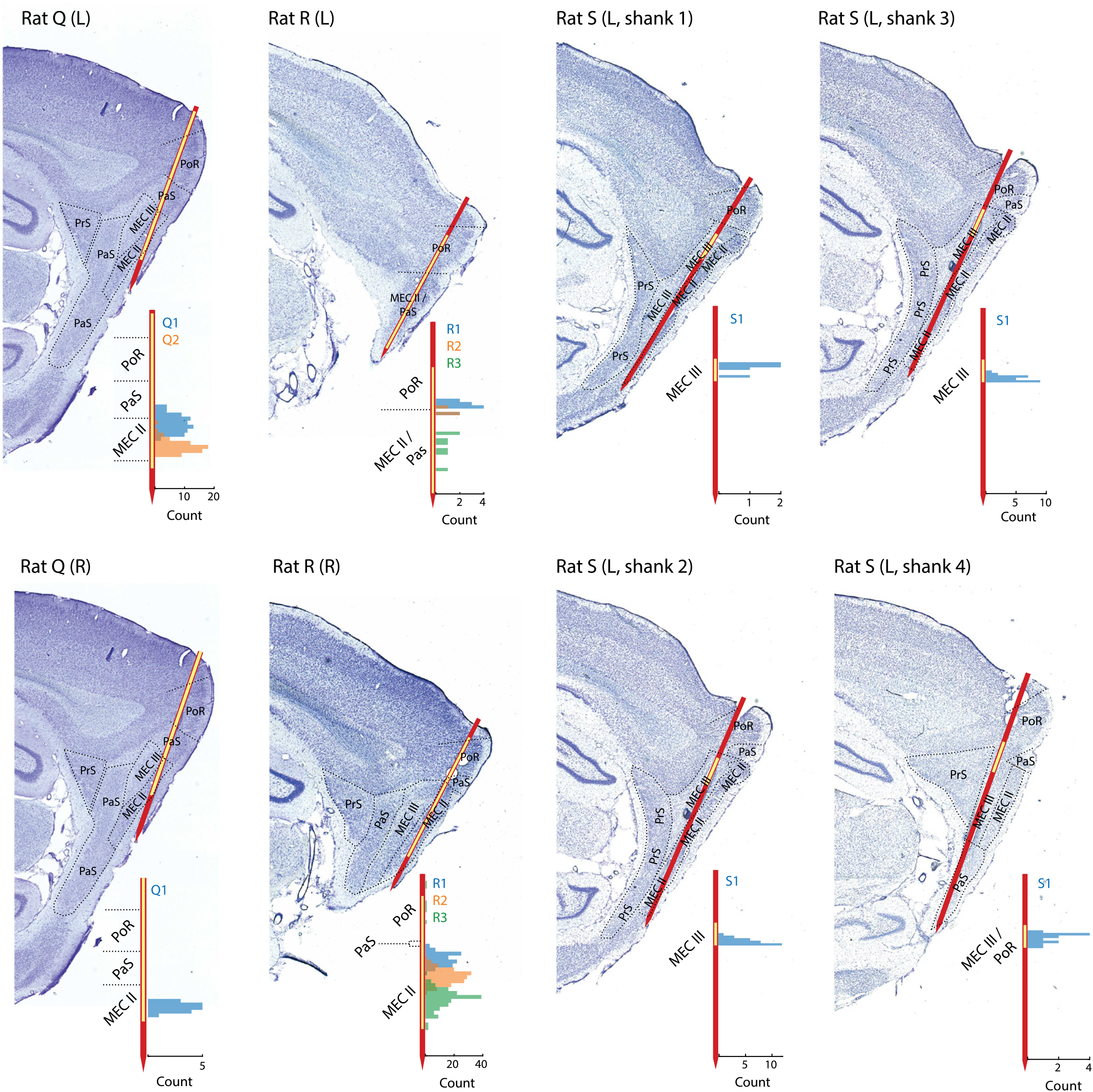
Nissl-stained sagittal brain sections showing recording locations for Rats Q, R and S. The probe shank is indicated in red (only the implanted portion of the shank is shown), with yellow overlay indicating the range of active recording sites. Stippled lines indicate borders between brain regions (MEC, medial entorhinal cortex; PaS, parasubiculum, PrS, presubiculum; PoR, postrhinal cortex). Layers are indicated for MEC (MECII, MECIII). Animal name, hemisphere (L, left; R, right) and shank number are indicated in text above each section. Insets show, for each section, the number of grid cells recorded at each depth of the electrode. Counts are colour-coded according to module identity. Note that several modules spanned across hemispheres.

**Extended Data Figure 2:**
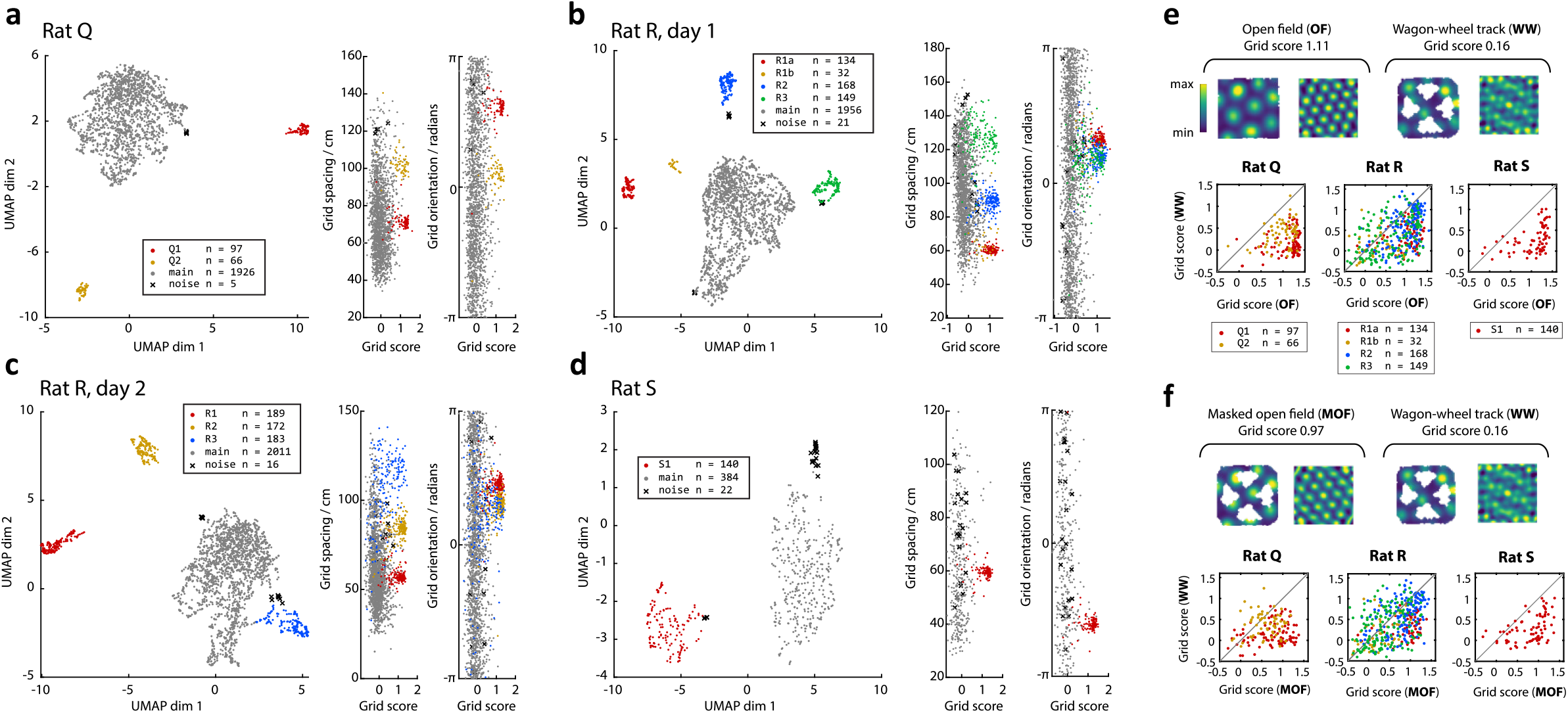
Grid module identification and properties. **a-d.** Clustering of grid modules (**a**, Rat Q; **b**, Rat R, day 1; **c**, Rat R, day 2; **d**, Rat S). For all experiments, coarse spatial autocorrelograms were first calculated from all cells’ open field firing rate maps (Extended Data Fig. 3). UMAP was then used to reduce the M-dimensional autocorrelograms (where *M* = 668 spatial bins) to a 2-dimensional point cloud, where each point represented the autocorrelogram of a single cell, and distances between points represented the similarity between autocorrelograms. Left scatterplot in **a**-**d**: 2D point cloud, with points colour-coded according to cluster ID. Clusters of cells corresponding to grid modules were identified by applying the density-based clustering algorithm DBSCAN to the 2-D point cloud. In every recording, the largest cluster (in grey, labelled “main”) comprised mainly non-grid cells, and the remaining smaller clusters (coloured) represented different modules of grid cells (1 module for Rat S, 2 modules for Rat Q, 3 modules for Rat R day 2; 4 modules for Rat R day 1). The DBSCAN clustering also identified outlier data points, shown here as black crosses and labelled “noise”. Right pair of scatterplots in **a**-**d:** Combinations of three grid parameters (grid score, grid spacing and grid orientation) for co-recorded cells from each recording. Each dot corresponds to one autocorrelogram (one cell). Dots are coloured by cluster ID as in **a**. The high grid scores of cells in the module clusters reflect the 6-fold symmetry of these cells’ grid patterns^55^. **e.** Comparison of grid-cell spatial periodicity in the open field arena (OF) and on the wagon-wheel track (WW). Top: firing rate map and corresponding autocorrelogram for an example grid cell in OF (left) and WW (right). For the purposes of this comparison, the same position bins were applied to both environments, resulting in cropping of the outermost parts of WW. Colour coding as indicated by scale bar; peak rates 16.1 Hz (OF) and 15.8 Hz (WW); range of autocorrelation values: −0.56 to 0.83 and −0.58 to 0.71, respectively. Note the more irregular appearance of the autocorrelogram for WW. Bottom: scatter plots showing grid scores of all grid cells in OF (x-axis) and WW (y-axis). Colours refer to the module assignment in **a**. Note the bias for points to lie in the lower-right quadrant, reflecting generally higher grid scores in OF than in WW. **f.** As for **e**, but controlling for differences in behavioural coverage of OF and WW environments. It is possible that the lower WW grid scores in **e** were a product of sparser behavioural coverage of the WW environment (animals visited only positions on the track). To control for this possibility, we created “masked OF” (MOF) rate maps by removing spatial bins from the original OF rate map which were not visited by the animal in WW. Top row shows the same example cell as in **e** after leaving a similar subset of position bins in OF as in WW. Bottom row shows comparison of grid scores for MOF and WW. As in **e**, grid scores are lower for WW, indicating that grid periodicity is reduced in WW even when differences in spatial coverage are accounted for.

**Extended Data Figure 3:**
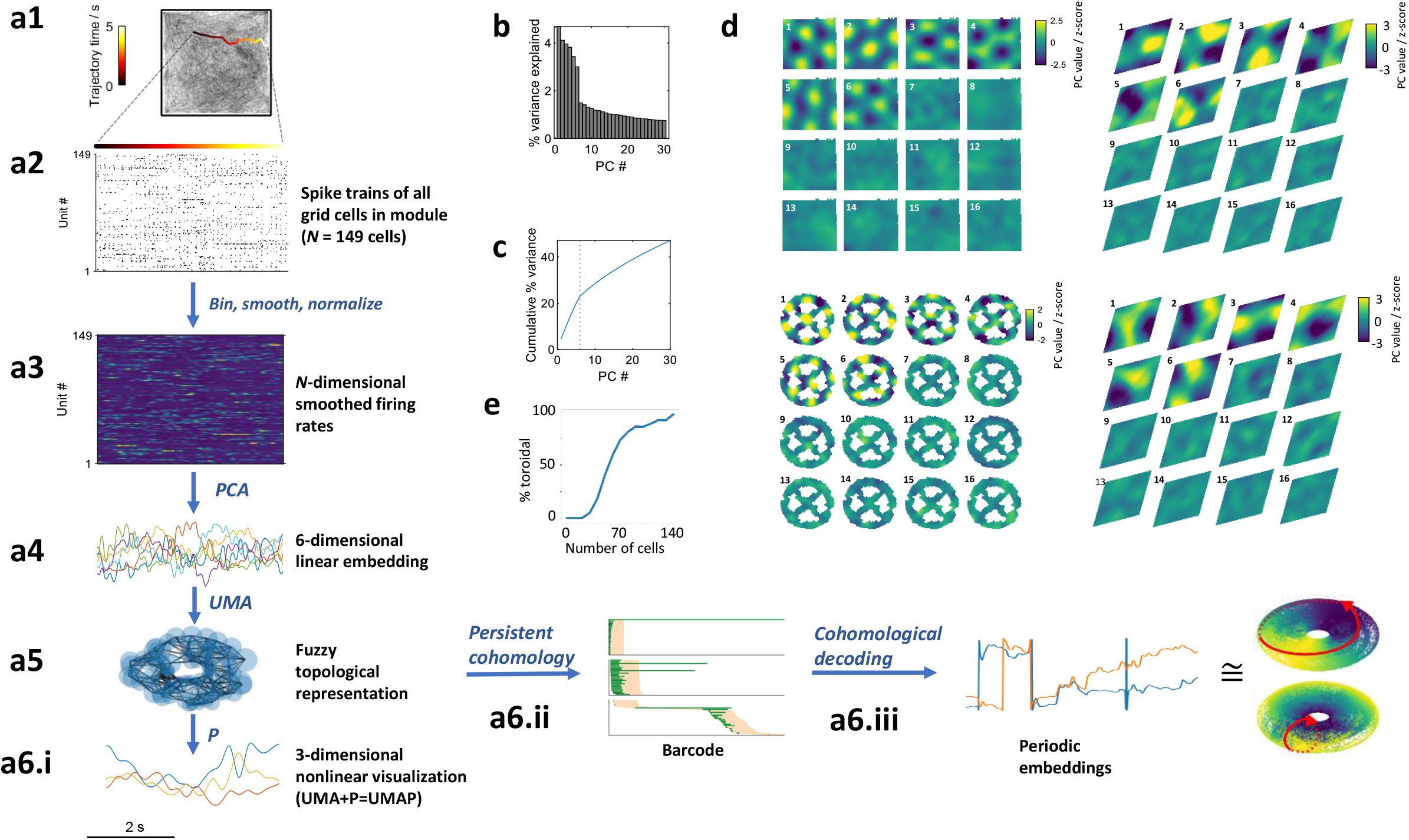
Preprocessing steps for visualization and detection of toroidal topology. **a.** Flow diagram showing method for extracting low-dimensional embeddings of neural activity. The animal forages in an open-field arena (**a1**), while spikes from 149 grid cells shown in Fig. 1a are recorded (**a2**; cells are ordered arbitrarily). A 5-second example behavioural trajectory is highlighted in **a1**, with colour indicating elapsed time. The spike trains are binned in time (*N* bins) and then smoothed and normalized, yielding a matrix of *N-*dimensional population activity vectors (**a3**). Linear dimensionality reduction (PCA) is applied to the matrix shown in **a3**, yielding a 6-dimensional linear embedding of the *N*-dimensional neural activity (**a4**). The six principal components are then passed through a second, nonlinear, dimensionality reduction step by UMAP, which generates a three-dimensional nonlinear embedding (**a6.i**) which allows the toroidal structure to be visualized. UMAP consists of two steps: first, a fuzzy topological graph representation is constructed (i.e. a “Uniform Manifold Approximation” - UMA) using a distance metric in the high-dimensional space (**a5**); second, to obtain the lower-dimensional projection (P), the coordinates of corresponding points in fewer dimensions are optimized to have a similar fuzzy topological representation. In the persistence analysis, we applied persistent cohomology to the fuzzy topological representation of the high-dimensional point cloud (**a6.ii**) and subsequently used cohomological decoding to obtain a two-dimensional projection of the original *N*-dimensional point cloud (left, showing a 5-second snippet). This may subsequently be visualized by using a three-dimensional toroidal projection of the two-dimensional angular values (right), as shown in Fig. 3a. The point cloud is coloured according to each angular coordinate, whose direction is indicated by a red arrow. **b.** Variance explained by the first 30 principal components (PCs) after applying PCA to the *N*-dimensional neural activity shown in **a3**. Note the particularly high values for the first six components. **c.** Same as **b** but showing cumulative explained variance. Note the pronounced ‘knee’ in the curve at the sixth PC. **d.** The first six PCs contain a population grid-like representation. Left: Each panel shows the mean value of one PC as a function of the animal’s position in the OF (top) or WW (bottom). PC value is colour-coded as indicated by the scale bar. PCs are ordered and numbered in red from top-left to bottom-right for each environment (OF, WW). Note the presence of grid-like structure in the first six PCs, which has a similar scale and orientation as individual cells in the grid module. These six grid-like PCs correspond to the set with the highest explained variance in **b**, each seemingly explaining a similar amount. Right (for OF and WW): Mapping the PCs onto the toroidal sheet given by the cohomological decoding (as in **a6.iii**), we see that each of these components completes one cycle in the fundamental tile. PC values are indicated by the scale bar. These components seem to reflect the proposition from CAN models that the isometric activity bump is translation-invariant with respect to the hexagonal symmetry of the neural sheet, which would suggest that the six first PCA components should correspond to the six fundamental Fourier modes of a hexagonal lattice (see Supplementary Methods for theoretical explanation of the six-dimensionality). **e.** The percentage of subsamples of R2 (resampled randomly 1000 times per number of cells) for which toroidal structure was detected in the parameterization given by the two most persistent 1-D bars in the barcode (as in Extended Data Fig. 4). Note that approximately 60 cells were needed for the probability of detecting toroidal structure exceed 50%.

**Extended Data Figure 4:**
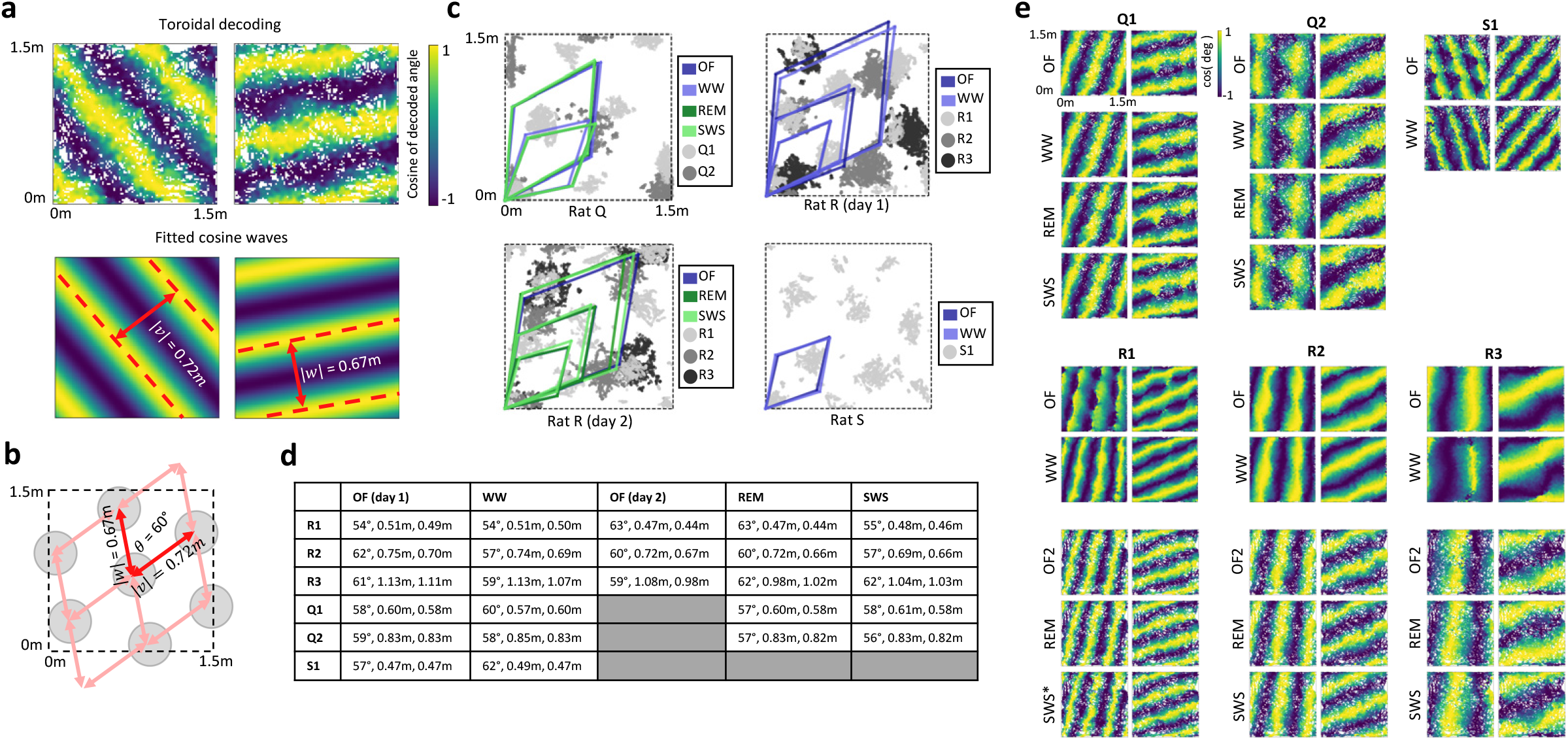
Mapping of decoded circular coordinates onto the open field allows geometrical interpretation of toroidal structure. **a.** Top row: Toroidal coordinates given by cohomological decoding from activity of grid module R2 during open field foraging, mapped onto the recording box. In each plot, colour indicates the mean value of the cosine of each of the two circular coordinates, computed in 100^2^ bins of physical space with spatial smoothing using a Gaussian kernel of standard deviation 1 bin. The mappings of both coordinates show 2-D striped patterns, with similar periods but distinct angles. Bottom row: A cosine wave is fitted to each coordinate to obtain the direction of the toroidal axes. The period and angle of the cosine wave in the plane may be represented by spatial vectors, *ν* and *w*, with corresponding length and orientation. Note the clear transversality of the two circles, expressed in the directions of the two vectors, further confirming the toroidal identification of the data. **b.** The periods and angles of the cosine waves in **a** reflect the scale and orientation of the grid module. Taking the origin of the vectors in **a** to be alike, we see that the vectors span a rhombus with approximately equal side lengths (0.67m and 0.72m) and an angle of 60 degrees. When repeated across the environment, the rhomboidal tile depicts the hexagonal grid pattern of the grid cell module, confirming that the product of the two decoded circles defines a hexagonal (“twisted”) torus. **c.** Rhombi of each module for each open field session, given by the cosine wave fitted to the toroidal coordinates (as in **b**). The toroidal parametrizations were obtained independently in different behavioural conditions (colour-coded), then used to decode the module’s activity during open field foraging, and subsequently mapped as a function of the position of the rat in the environment (see **e**). Positions of downsampled spikes from example cells of each module are shown in greyscale to illustrate grid scale and orientation. The consistent angle and side lengths suggest the geometry of the rhombus is retained across brain states and environments, with a constant scale relationship between modules. **d.** Table of side lengths and angles of the cosine waves that form the rhombi in **c**, shown for each grid module and each condition. **e.** Visualization of the cohomological decoding of toroidal coordinates as a function of physical space (one visualization for each grid module during each condition, with the toroidal parametrizations aligned to common axes before creating the rate maps. All data sets where the barcodes indicated toroidal structure exhibited periodic stripes in the open field, with phase and orientation corresponding to the two-dimensional periodicity of the grid pattern of the respective module. SWS* refers to the decoding when considering only Class C_1_ of R1 as given by the correlation clustering method described in Fig 5b.

**Extended Data Figure 5:**
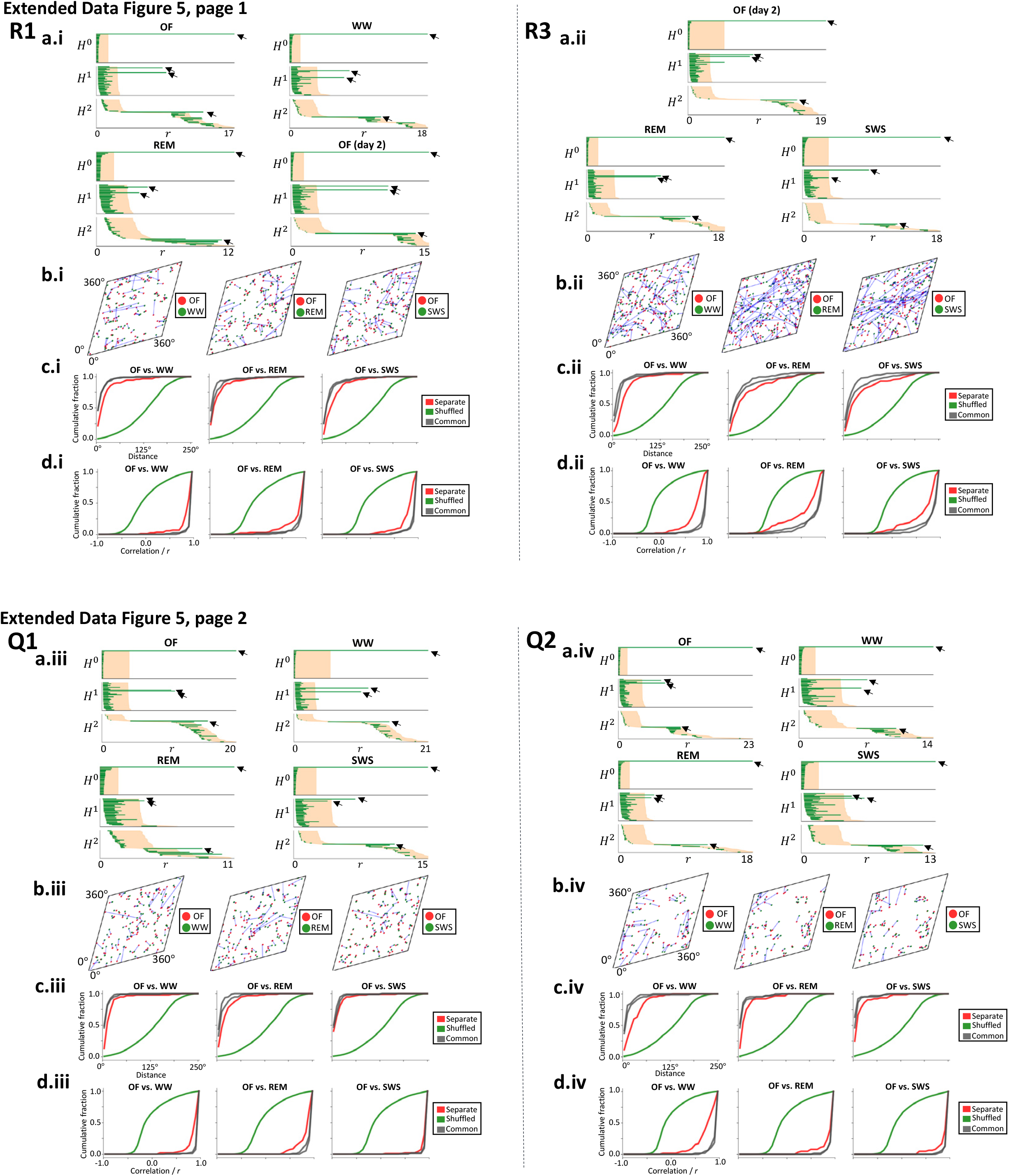
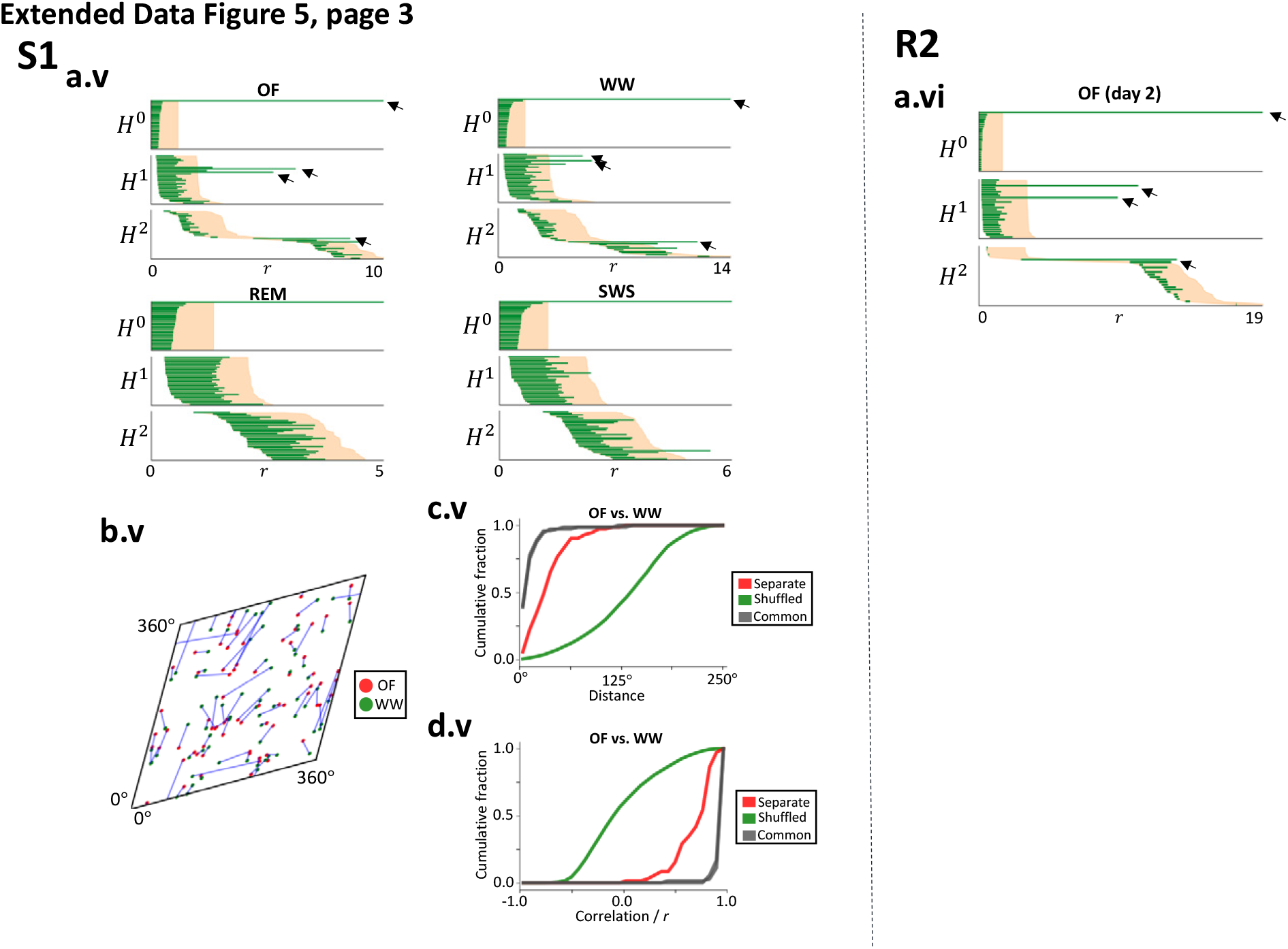
Barcodes and toroidal tuning statistics for grid modules or recording sessions not included in Fig. 2–5. Data are shown for six grid cell modules: R1, R3, Q1, Q2, S1 and R2. Toroidal structure is clearly present across environments and behavioural states. Captions for successive panels on each page: **a. i-vi** Barcode diagrams (as in Fig 2c, d) showing the results of the persistent cohomology analysis on open-field (OF), wagon-wheel track (WW) or sleep (REM or SWS) data. Arrows point to the longest-lived bars equivalent to those of a torus (one 0-D bar, two 1-D bars and one 2-D bar). **b. i-v** Distribution of grid cells’ receptive field centres on the inferred torus for OF and WW as well as sleep states, similar to Fig 3e. Each dot signifies the field centre of an individual grid cell. Blue lines connect field centres of the same cell across conditions. Note the proximity of red-green pairs (after separate alignment for the two recording sessions of each panel). **c. d.i-v** Cumulative distributions showing stability of grid cells’ toroidal tuning between brain states, as in Fig. 3f, g. Distributions show peak field distance (**c**) and Pearson correlation of pairs of toroidal rate maps (**d**). Red line: recorded data, decoding from toroidal representation in same environment (“Separate”, as in **b**); green line: shuffled data; grey lines: recorded data, same as for red line but using a single toroidal parametrization (“Common”, either OF, WW or sleep state) to decode the activity in both conditions. The strong displacement of the distributions for shuffled data relative to the distributions for original data indicates that the preservation of toroidal selectivity between OF and WW, OF and REM, and OF and SWS is far beyond chance level.

**Extended Data Figure 6:**
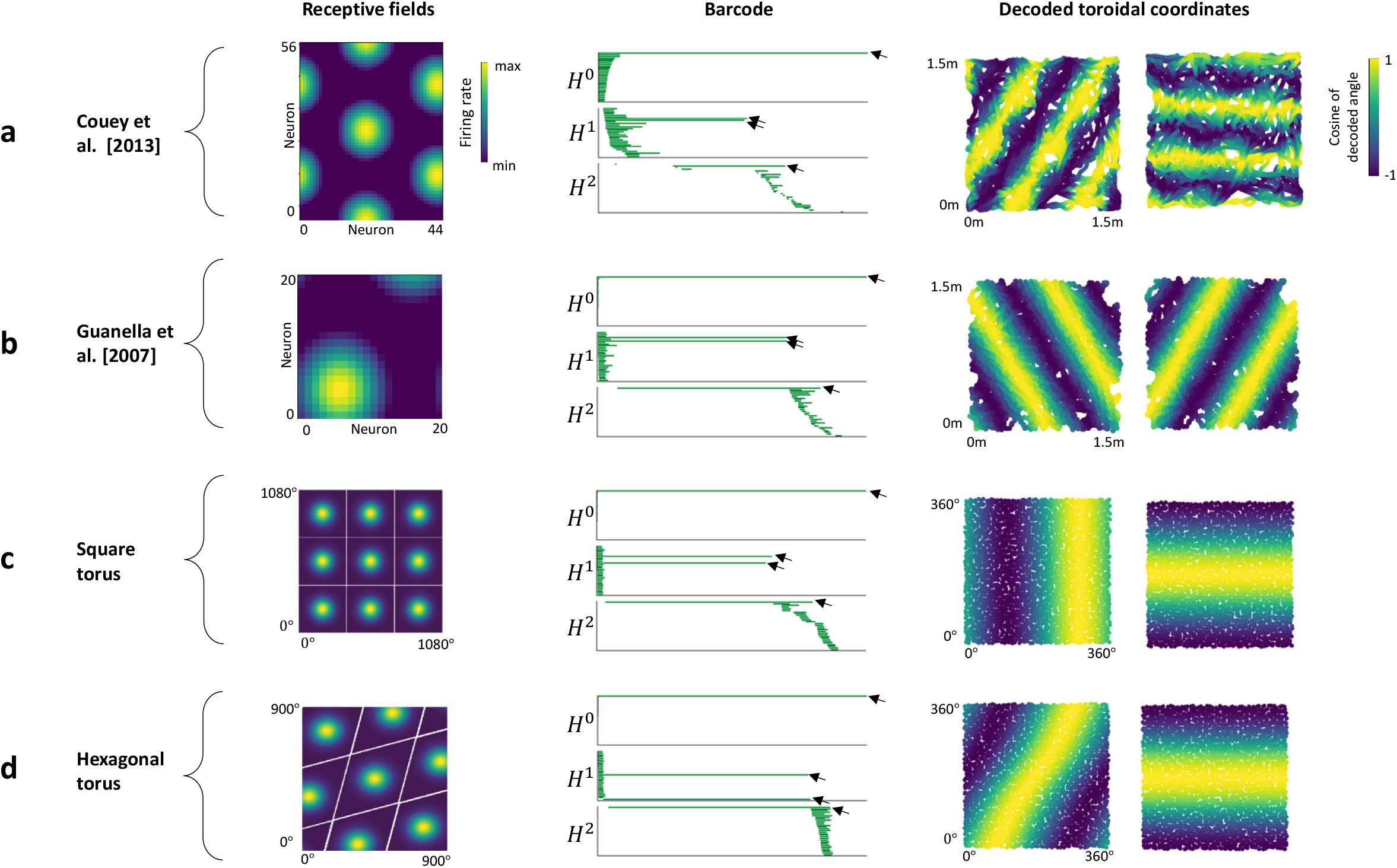
Barcodes and decoding of simulated firing activity for two grid cell CAN models (with no noise), and for two point clouds randomly sampled on a hexagonal and a square torus. **a.** Persistent cohomology analysis of a simulated grid-cell network based on the CAN model from Couey et al (2013)^15^ during open-field foraging (using the position coordinates from 1000 seconds of the open field session of rat ‘R’, day 1). Left: Colour-coded firing rates for a single time frame of the 56 × 44 grid cells, shown at their respective positions on the neural sheet. Note the multiple bumps of activity. Middle: Barcode of the simulated data. Arrows point to one 0-D, two 1-D and one 2-D bar with long lifetimes, indicating toroidal structure. Note the similarity with the barcodes of the original data (Fig 2c-f, 4, Extended Data Fig. 5a). Right: Each coordinate of the toroidal parametrization of the two longest lived 1-D features is mapped onto the spatial trajectory, colour-coded by its cosine value (as in Extended Data Fig. 4a, e). The resulting striped patterns of the two maps are oriented approximately 60 degrees relative to each other, as expected from a hexagonal torus network structure (see **d**). **b.** Analysis of a random sample of 100 grid cells from a 20×20 simulated grid cell network, using the twisted torus CAN model formulated by Guanella et al (2007)^16^ during 5000 time frames of a simulated random open field walk. Left: Firing rates of the cells in the network at a single time frame. The model generates a single bump of activity based on both inhibitory and excitatory, asymmetric connections representing a twisted torus. Barcode (middle) and cohomological decoding of toroidal position (right) are shown as in **a**. The barcode shows four prominent bars: one 0-D bar, two 1-D bars and one 2-D bar, similar to that of a torus. Note that the pair of stripes in toroidal coordinates are oriented 60 degrees relative to each other. **c., d.** To verify the expected barcodes and decoding of a torus and compare with both real and synthetic grid cell data, we performed the same topological analysis on point clouds sampled from two idealized toroidal parametrizations: a 4D description of a square torus (**c**) and a 6D _embedding_ of a hexagonal torus (**d**). Left: Representing the firing of a cell as a Gaussian function centred at a single toroidal coordinate on the toroidal sheet results in a square (**c**) and hexagonal (**d**) firing pattern, when arranged to tesselate a 2-D surface. Middle: The expected barcode of a torus (one 0-D, two 1-D, and one 2-D bar clearly longer than the other bars) is seen in both cases (top). Note the cluster of short-lived 1-D bars born shortly after the births of the two significant 1-D features and the cluster of short-lived 2-D bars born shortly after the deaths of these long-lived features. Although the barcodes cannot distinguish the geometries of the tori, when mapping the decoded toroidal coordinates onto the sampled angles (bottom), we note a clear difference between the stripe patterns of the two tori. The decoding of the square torus displays orthogonal stripes, in contrast to the 60° angle of the stripes of the decoded hexagonal torus; thus, only the latter resembles our findings in the recorded and simulated data.

**Extended Data Figure 7:**
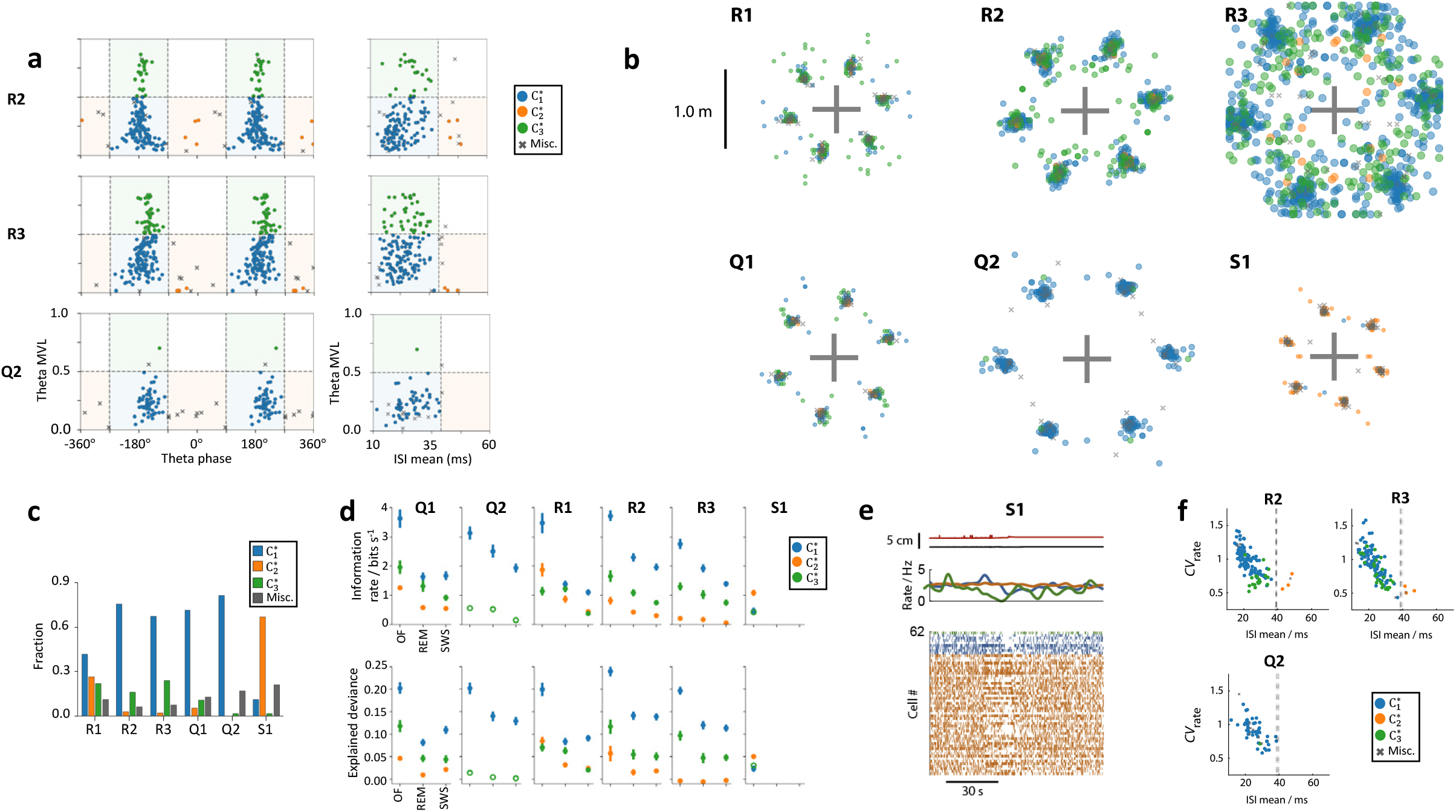
Subpopulations of grid cells with different temporal spiking statistics have different degrees of toroidal selectivity. **a.** Grid cells of the same module segregate into functional classes with distinct spike timing characteristics (as shown for R1, Q1 and S1 in Fig. 5e). Plots are as described for Fig. 5e, but showing the three remaining modules (R2, R3 and Q2). **b.** Geometry of grid-cell pattern of all six modules with classes. Each plot shows the locations of the innermost six peaks of the spatial autocorrelogram for every grid cell in one module. Each dot indicates the position of one peak from one cell (total of 6 dots per cell); dots are coloured by the cell’s class 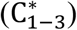, as indicated in box to the left. The grey crosshair indicates the center of the autocorrelogram. **c.** Frequency distribution showing fraction of 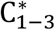 cells belonging to each class after applying the classification in Fig. 5e on all 6 grid modules. Modules R1 and S1 stand out with a larger proportion of cells in 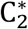. **d.** Mean toroidal information rate and explained deviance (± S.E.M.) for each class during OF, REM and SWS. Points are color-coded by class 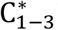. 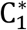 shows higher mean information rate and explained deviance scores than both 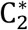 and 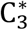 in all brain states, except for S1 which does not display toroidal invariance across brain states. Note only 1 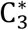 cell for S1 and Q2 (open circles). **e.** Low-activity states during rest. Similar to Fig. 5f but for module S1. Note that although the average firing rate of 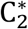 cells is not strongly perturbed by the low-activity state, individual cells show sustained increases or decreases in firing rate. Similar low-activity states have been described previously in MEC and other forebrain regions^68,69^. **f.** Scatterplots showing that bursting grid cells (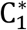 and 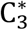) express more slow-timescale firing rate variability during SWS. Plots are as for Fig. 5g but show the three other modules.

**Extended Data Figure 8:**
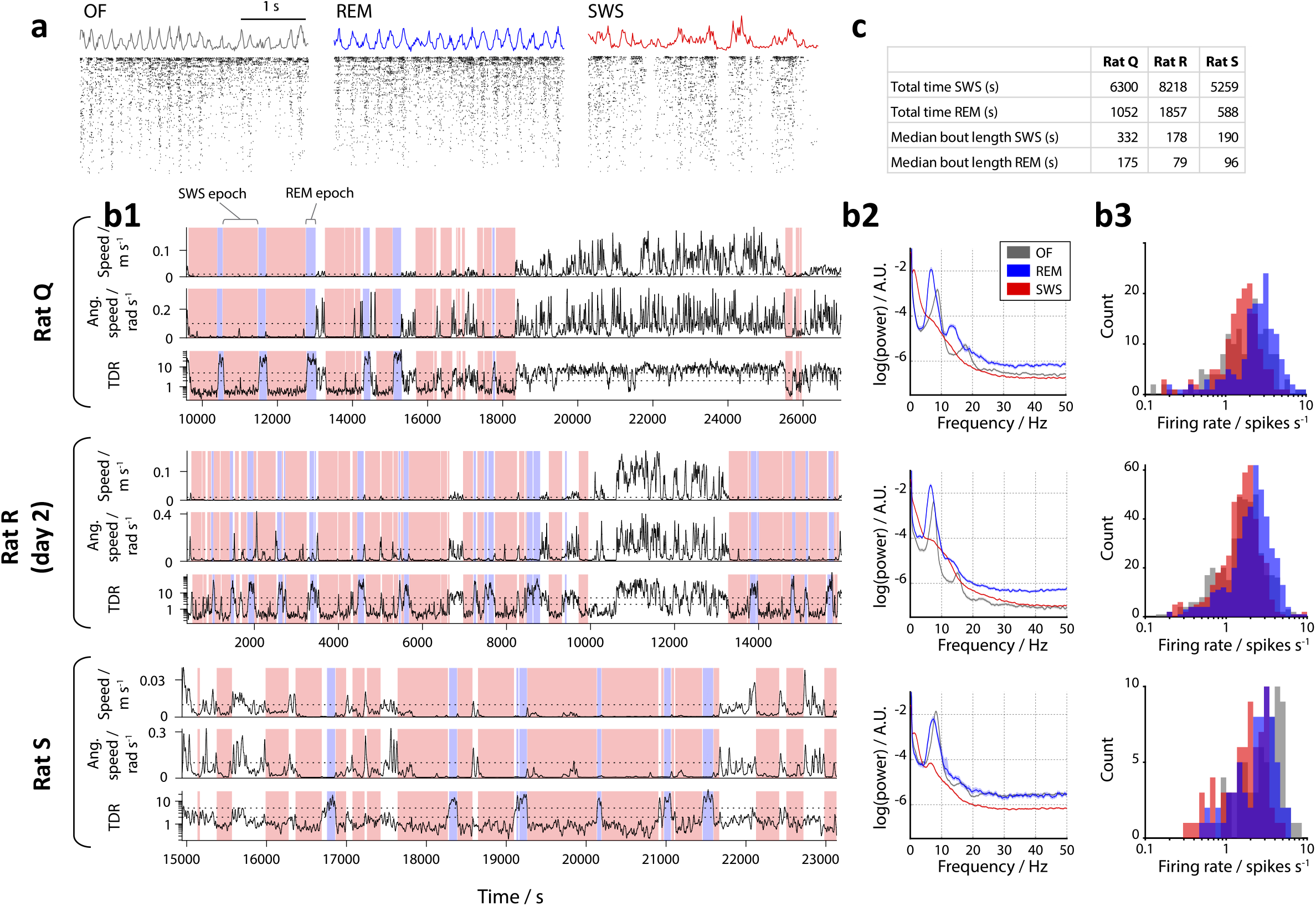
MEC spiking activity across different sleep/wake states. **a.** Example traces of MEC multi-unit activity (upper; coloured lines), and rasters of spike times of 444 grid cells (lower; black dots) recorded from rat R during during open field foraging (OF), REM sleep and slow-wave sleep (SWS). Cells are ranked from top to bottom by the number of spikes fired during the example time window. Note the presence of regular theta waves (5—10 Hz) during OF and REM, and presence of slower, more irregular fluctuations and silent “down-states” during SWS. **b.** Classification of sleep/wake states based on behavioural and neural activity during rest sessions. Each of the three vertical blocks shows a recording from one animal. Rat R day 2 did not contain a rest session and is not shown on this figure. **b1:** Detection of REM and SWS sleep epochs in the rest session. The plots show the time courses of the three variables used for detecting REM and SWS epochs. Top panel of each block: animal locomotion speed; middle panel: the animal’s head angular speed; bottom panel: the ratio of the amplitude of theta (5— 10 Hz) and delta (1–4 Hz) frequency bands in the multi-unit spiking activity (theta/delta ratio, TDR). Sleep periods were defined by sustained immobility (> 120 s with locomotion speed < 1 cm s^-1^ and head rotational speed < 6° s^-1^). These sleep periods were subclassified into SWS (red shading) and REM (blue shading) based on TDR: periods where TDR remained above 5.0 for at least 20 s were classified as REM; periods where TDR remained below 2.0 for at least 20 s were classified as SWS. **b2.** Log-power spectra of MEC multi-unit activity during each sleep/wake state. The power spectra were calculated in 5-second windows using the multitaper method (see Online methods), and using all available time periods for each state. The line and shaded area indicate the mean and 95% bootstrap confidence intervals, calculated across time windows (confidence intervals are narrow). Note the pronounced peak corresponding to the theta band (5–10 Hz) during OF and REM, and the higher power in the delta band (1–4 Hz) during SWS. **b3.** Histograms showing distributions of firing rates for all grid cells during each sleep/wake state (number of grid cells: rat Q 159, rat R 428, rat S 72). **c.** Table showing total time and median bout length of recorded sleep for each animal.

**Extended Data Figure 9: Page 1-9.**
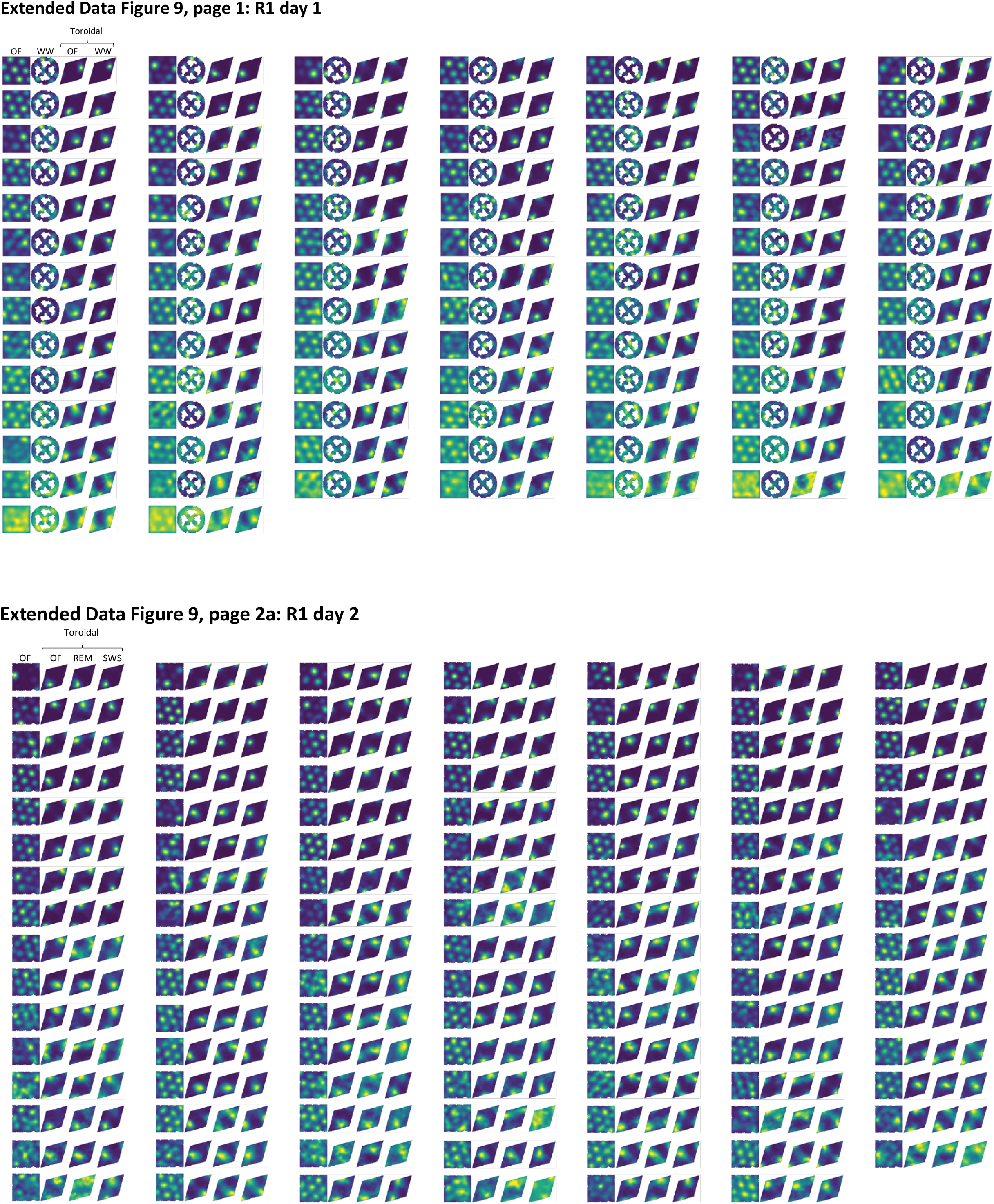
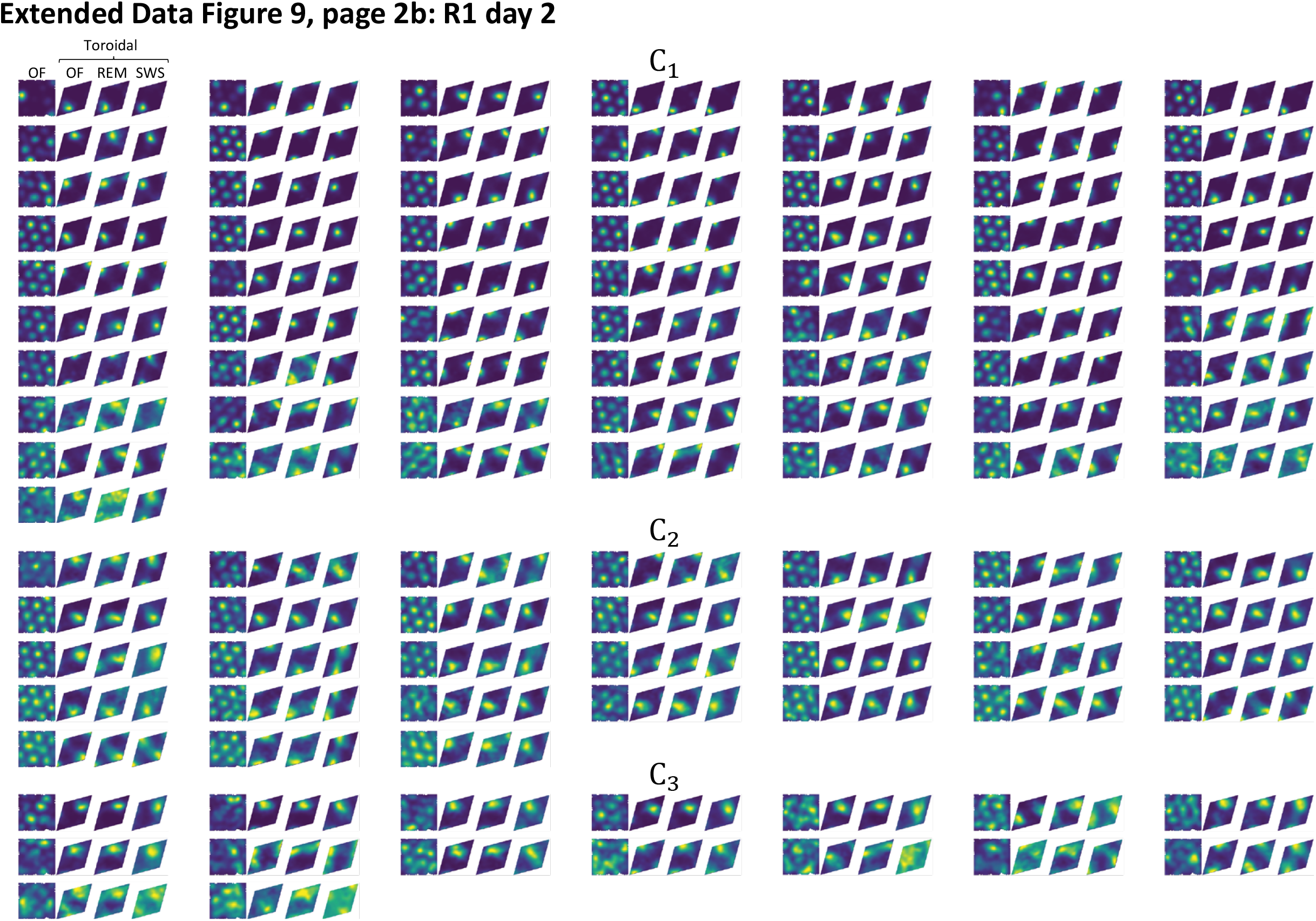
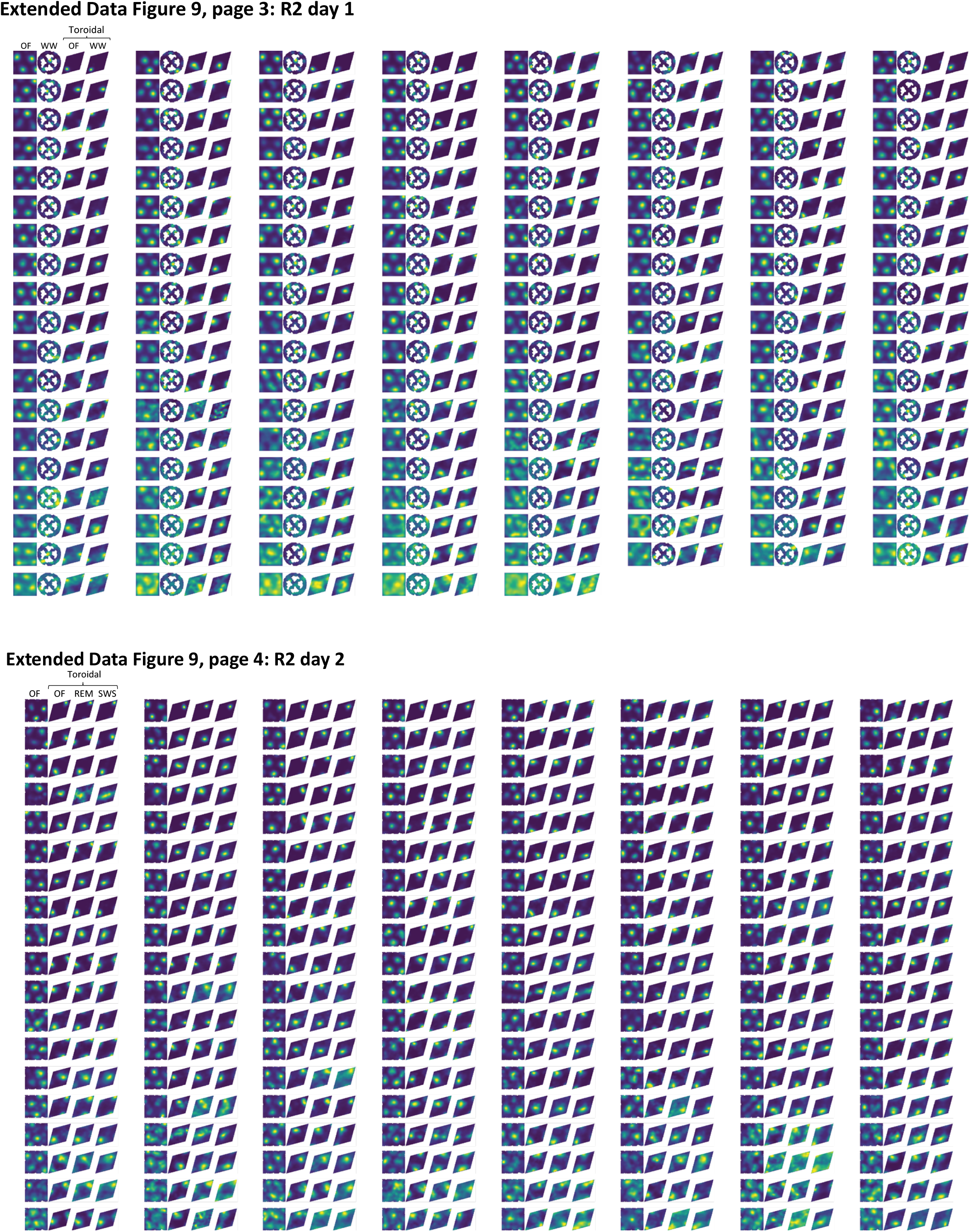
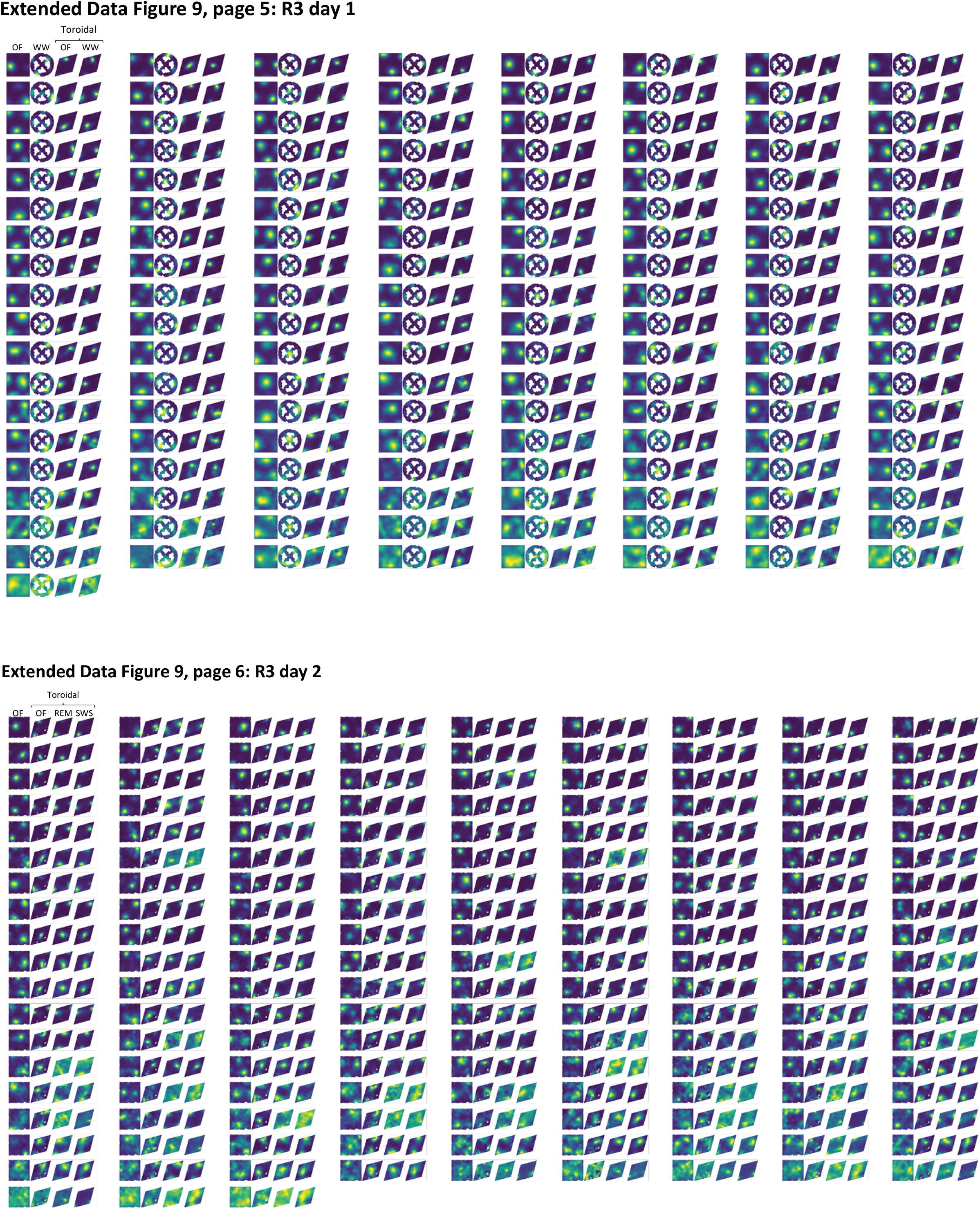
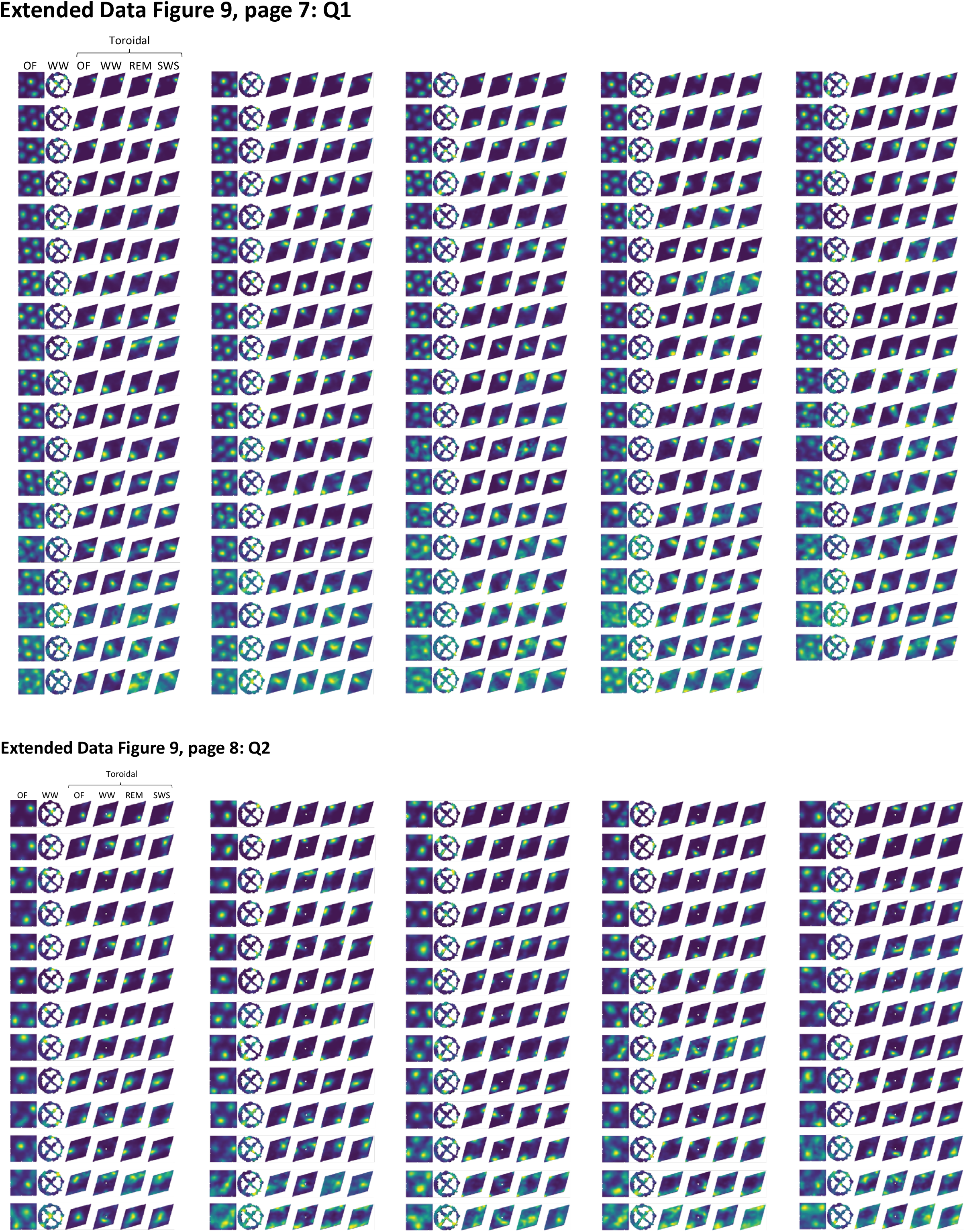
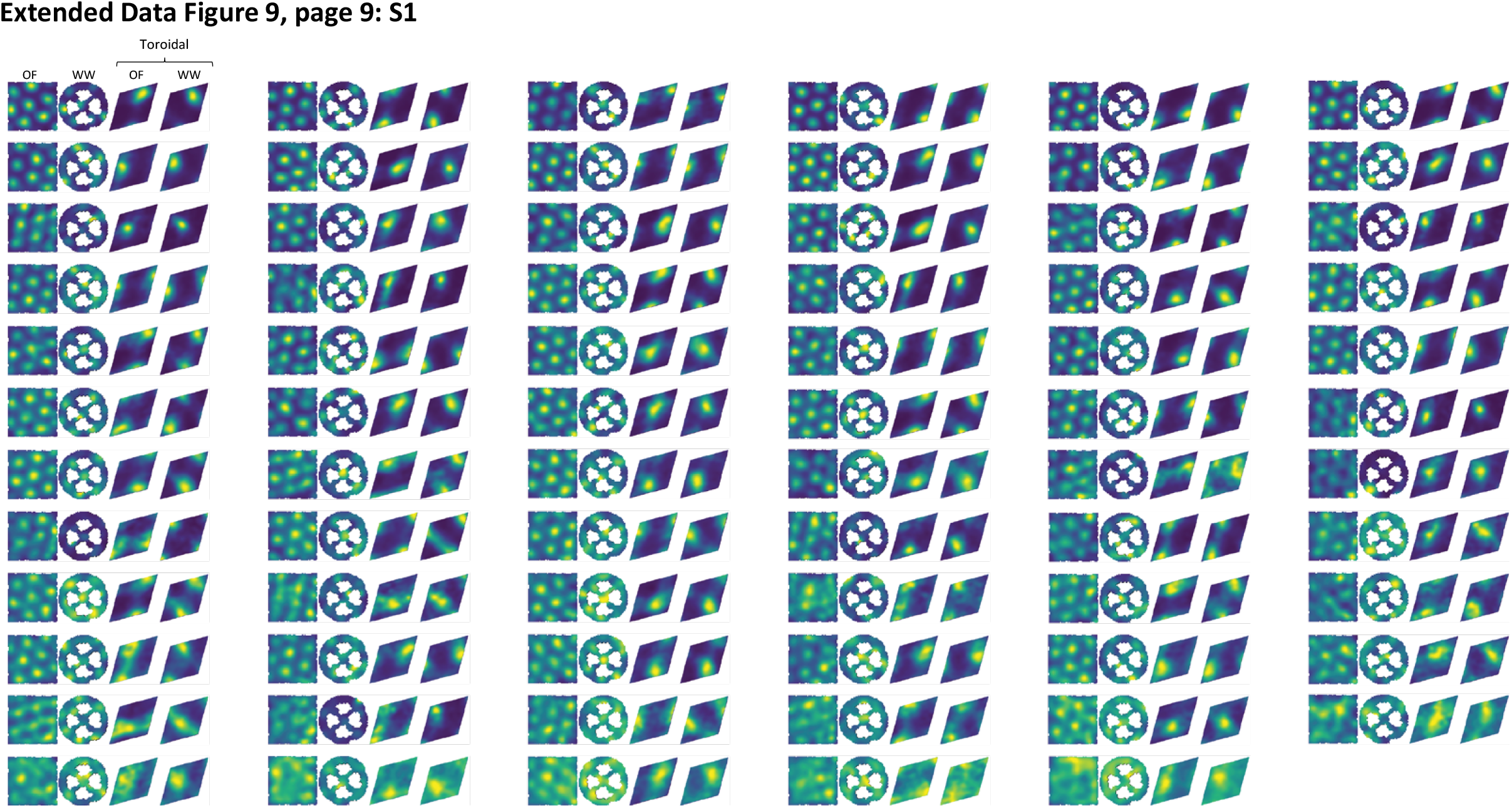
Each page shows the tuning, for all grid cells from one module in one recording, to coordinates on the inferred torus. Subsets of these plots are shown in Fig. 3b, 4b, 5d. Rate maps show the normalized firing rate of individual cells as a function of position in the spatial environment (open field OF and wagon-wheel track WW; leftmost columns) or on the inferred torus (for OF, WW, REM and/or SWS; rightmost columns). Page 2b shows the same plots as Page 2a, but with the cells subdivided into their respective classes (C_1-3_; see Fig. 5). Note that relative firing locations on the inferred torus are consistently preserved between environments and behavioural states.

## Acknowledgments

We thank M.P. Witter for help with evaluation of recording locations, and A.M. Amundsgård, K. Haugen, K. Jenssen, E. Kråkvik, Ingvild Ulsaker-Janke, and H. Waade for technical assistance. The work was supported by a Synergy Grants to E.I.M. and Y.B. from the European Research Council (‘KILONEURONS’, Grant Agreement N° 951319), an RCN FRIPRO grant to E.I.M. (grant number 286225), a Centre of Excellence scheme grant to M.-B.M. and E.I.M. and a National Infrastructure grant to E.I.M. and M.-B.M. from the Research Council of Norway (Centre of Neural Computation, grant number 223262; NORBRAIN, grant number 295721), the Kavli Foundation (M.-B.M. and E.I.M.), the Department of Mathematical Sciences at the Norwegian University of Science and Technology (B.D., E.H., N.A.B.), a direct contribution to M.-B.M. and E.I.M. from the Ministry of Education and Research of Norway, and grants to Y.B. from the Israel Science Foundation (grant No. 1745/18) and the Gatsby Charitable Foundation. Some of the computations were performed on resources provided by the NTNU IDUN/EPIC computing cluster.

## Author Contributions

R.J.G., M.-B.M. and E.I.M. designed experiments; R.J.G. performed experiments; N.A.B., E.H., B.A.D., R.J.G., Y.B. and E.I.M. conceptualized and proposed analyses; E.H. and R.J.G. developed and performed the analyses; M.P. shared unpublished Kilosort software; R.J.G., E.H., B.A.D., Y.B., M.-B.M. and E.I.M. interpreted data; E.H. and R.J.G. visualized data; R.J.G., E.H., B.A.D., Y.B. and E.I.M. wrote the paper, with periodic input from all authors; E.I.M., M.-B.M., B.A.D. and N.A.B. supervised the project; E.I.M., M.-B.M. and Y.B. obtained funding.

## Supplementary Information

is available for this paper.

## Author Contact Information

Correspondence should be addressed to E.I.M., R.J.G., N. A.B. or B.A.D. Requests for materials should be directed to E.I.M.

## Reprints and Permissions

information is available at www.nature.com/reprints

## Supplementary Methods

### Theoretical explanation of the six-dimensionality proposed by PCA

To understand why a minimum of six PCA components is necessary in order to account for a large fraction of the variance in the population patterns of grid cells, we shall consider an idealized model of grid cell firing, with the following three assumptions:

First, grid cell population activity patterns lie on a two-dimensional manifold with toroidal topology. This 2-D surface can be mapped to a rhombus with an angle of 60 degrees and periodic boundaries. Specifically, we can parametrize positions on the rhombus in the form 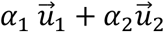, where 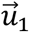, and 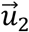 are unit vectors whose orientations differ by 60 degrees, and α_1,2_ ∈ [0,1].

Second, the tuning of an individual grid cell to the toroidal coordinates (the position on the rhombus) is identical in all cells, up to a translation:

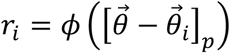

where 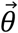 is the 2-D position on the rhombus, 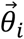 is the center of the receptive field of cell *i*, 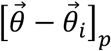 represents a shift of 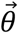 by 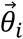 with periodic boundary conditions on the rhombus and *φ* represents the structure of the tuning function.

Third, activity patterns of the neural population uniformly sample the states that correspond to different positions on the rhombus.

Under these conditions, the covariance matrix *C* of neural activity has the structure

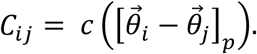

If we further assume that receptive field centers 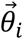 uniformly and regularly sample the rhombus, the covariance matrix commutes with periodic rigid translation operators on the rhombus. A full basis of eigenvectors of *C* can then be obtained such that the eigenvectors are also eigenvectors of the translation operators. Thus, these eigenvectors are Fourier modes of the form

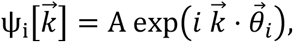

where the wavevector 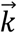 must be selected such that *ψ* is periodic on the rhombus. To achieve this requirement, 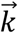 must be a vector lying on a vertex of a triangular lattice: the reciprocal of the lattice with basis vectors 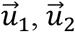.

The eigenvalues of 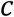 are the corresponding Fourier transform components of *c*. Typically, for a unimodal tuning curve, these eigenvalues will be a monotonically decreasing function of 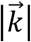 for 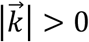. Furthermore, if the tuning of individual cells is isotropic, the eigenvalues depend only on 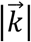. Therefore, the six PCA modes that correspond to the smallest value of 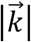 must contribute equally to the variance. If the tuning curve of individual cells is sufficiently wide, the eigenvalues are expected to decay rapidly with 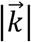, and the modes that correspond to the first six PCA components are expected to capture a large fraction of the variance.

### Persistent (co)homology

As persistent cohomology is a cornerstone of the current analyses, we wish to elaborate on its theoretical background to elucidate how and why it works (broader perspectives on topological data analysis and its application in biology are available elsewhere^87,88,95^). We start by introducing (co)homology and the Vietoris-Rips complex before we turn to persistent homology.

Given a topological space *X*, we can assign, for all natural numbers *n*, a vector space *H_i_*(*X*), called the *i*-th homology group of *X*, such that if *f: X → Y* is a continuous map between topological spaces *X* and *Y*, then *f_*_*: *H_i_*(*X*) → *H_i_*(*Y*) is a linear map between vector spaces. Similarly, we may define cohomology groups, *H^i^* (*X*) by reversing arrows. These are dual notions and give the same results in our case. We will thus only continue describing the former. The dimension of *H_i_*,(*X*) is called the *i*-th Betti number, representing the number of *i*-dimensional holes (for further details see Hatcher, 2002^89^). However, this may vary depending on the choice of coefficients of the vector space (note that it is choosing algebraic fields as coefficients that makes the homology groups vector spaces). For example, using ℤ_2-_ coefficients, the Klein bottle will have the same homology as a torus. To separate these, we use ℤ_47_-coefficients in our computations. The choice of field coefficients simplifies computations at the risk of losing topological information known as *torsion*, measuring the orientability of a space. However, the same number of holes (Betti numbers) is computed, which is what is here used to distinguish spaces.

As a point cloud is finite and discrete, its homology only returns the number of points in the point cloud (its 0-th Betti number). Thus, we associate combinatorial spaces known as simplicial complexes to the point cloud which may have non-trivial topology reflecting interesting structure and information of the data set and whose homology is easy to compute. A simplicial complex is a set *V* of vertices and a set S of finite non-empty subsets of *V* called simplices such that any vertex is a simplex and any non-empty subset of a simplex is a simplex. A simplex of cardinality *p* + 1 is referred to as a p-simplex (of simplicial dimension *p*) and geometrically, we may refer to a 0-simplex as a point, a 1-simplex as an edge, a 2-simplex a triangle, a 3-simplex a tetrahedron and so on in higher dimensions.

There are different choices in constructing simplicial complexes associated with the data. We used what is known as the Vietoris-Rips complex, here denoted *R_r_*. The vertices of the Vietoris-Rips complex are the points in the point cloud and the simplices are the sets of points whose pairwise distance is less than the scale value, r. This is equivalent to replacing each point by a ball of common radius *r* and connecting two points with an edge if their balls intersect. p-simplices are then formed if each point of a subset of *p* + 1 points have edges to all other points in the subset (i.e. a *p* + 1-clique). Although this simplicial complex is not homotopy equivalent to taking the union of balls (as e.g. the Čech complex is) and thus does not necessarily have the same homology, the basic topological information is preserved under this correspondence.

One way to construct the Vietoris-Rips complex in detecting the topology of neural data is to regard individual cells as points and their pairwise dissimilarity (e.g. correlation) as scale. In our case, we rather considered the population activity vectors as the points. There is a subtle correspondence between these constructions, where the resulting barcodes are the same when applying persistent cohomology. This relationship is only valid when the tuning of the cells is such that the response is convex, seemingly invalidated by the firing patterns of grid cells with respect to its physical position in the environment. However, when considering the tuning to be a function of the toroidal state space, we find it indeed to be convex (indicated by the single bumps in the toroidal rate maps for each grid cell – Fig 3a, Extended Data Fig. 9)^59^, suggesting both constructions should give rise to the same barcodes.

We consider the nested chain of Vietoris-Rips complexes, *R*, constructed for all increasing values of *r* in which new simplices are formed:

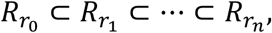

where *r_0_* = 0 and *r_n_* is the largest pairwise distance in the point cloud, and apply homology to get a sequence of vector spaces and maps, for all dimensions *i:*

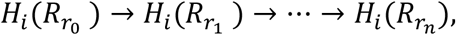

where the maps are induced by the inclusion maps (note that we have omitted compositions and identity maps), called the *i*-th persistent homology. This may again be decomposed into a sum of elementary *persistence modules^96^:*

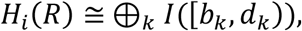

where *b_k_* < *d_k_* give the scales in which a class in *H_i_*(*R*) first appears and later disappears. Thus, we may represent the persistent (co)homology by displaying the intervals as bars starting at *b_k_* and ending at *d_k_*. The collection of bars for all dimensions results in what is known as the *barcode*.

The barcodes are shown to be stable under influence of noise (given certain assumptions on the construction)^97^. This means that small perturbations to the point cloud lead to small changes in the barcode. Thus, we note that in dimensionality reduction, quantifying the dissimilarity between the barcode of the high-dimensional representation and its embedding may address the challenge of measuring how faithful an embedding is^57^.

### The Cup Product

The barcode is only based on the additive structure of the cohomology vector spaces, which means that, for instance, the barcode of the one-point union of a sphere and two circles is equivalent to that of a torus. Introducing a multiplicative structure of cohomology – the *cup product* – helped confirm the toroidal structure.

To compute the cup product structure of the point cloud, we chose a scale that defined the Vietoris-Rips complex in which the significant features identified by the barcode (representative of a torus) existed. Due to computational reasons, we had to down-sample the point cloud to 300 points, for which only the population activity of the open field session for grid cell module R2 still showed clear toroidal structure. Doing so for this data set resulted in non-trivial cup product structure in dimension one, eliminating the possibility of a two-sphere with two circles attached, and strengthening the conjecture of toroidal topology of the underlying manifold of the grid cell population activity. For this computation we used the software *ChainCon^98^*.

The multiplicative structure may also be seen via the circular parametrizations of the point cloud. Mapping the two circular features given by the barcode onto the open field arena (Extended Data Fig. 4), we see that the generators (that give rise to the circular maps) correspond to cycles intersecting transversally. Hence, these should have non-trivial cup product, and thus clearly indicate a toroidal state space. We note, however, that a subtler description of the topology of the state space would lie in the *homotopy type* of the space^89^.

---

**Supplementary movie 1:** 3-D UMAP visualization of toroidal manifold.

The 3-D point cloud shows a UMAP embedding of the activity of 149 grid cells from module R2 in the open field arena, as shown in Fig. 1b, c, d. Each dot represents the population activity state at one point in time. Dots are coloured by the value of the first principal component of the population activity.

**Supplementary Figure 1 for Reviewers:**
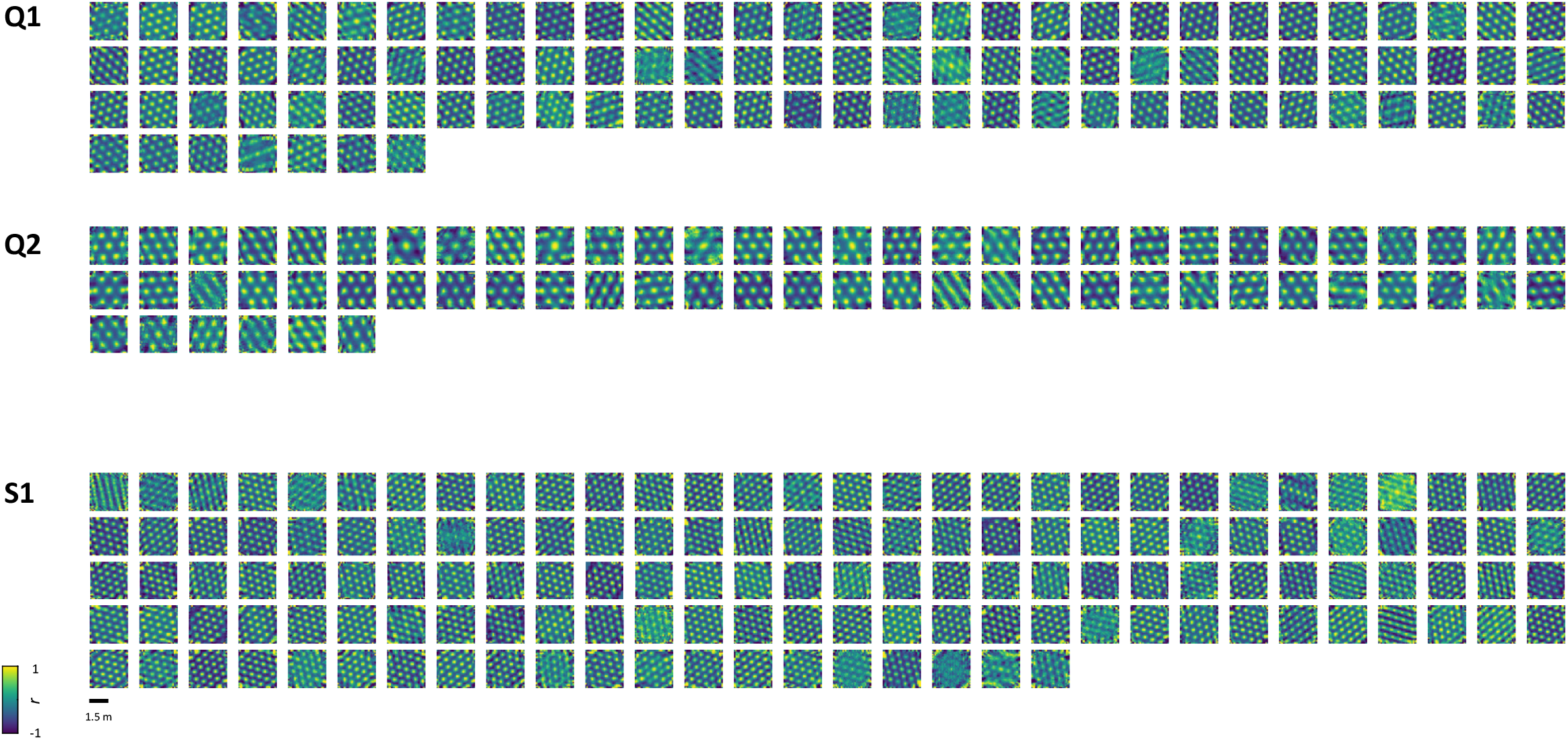
Coarse spatial autocorrelograms of all grid cells. Each plot shows the coarse 2-D spatial autocorrelogram generated from the firing rate map of one cell during open field foraging (total of 97 cells for module Q1; 66 cells for Q2; 140 cells for S1). The rate map used to compute each autocorrelogram was constructed using a 15×15 grid of 10-cm bins. Autocorrelation values range from −1 to 1 (colour scale bar). Scale bar for box size, 1.5 m. Plots (cells) are grouped by grid module membership as assigned by the clustering shown in Extended fig. 2a-d.

**Supplementary Figure 2 for Reviewers:**
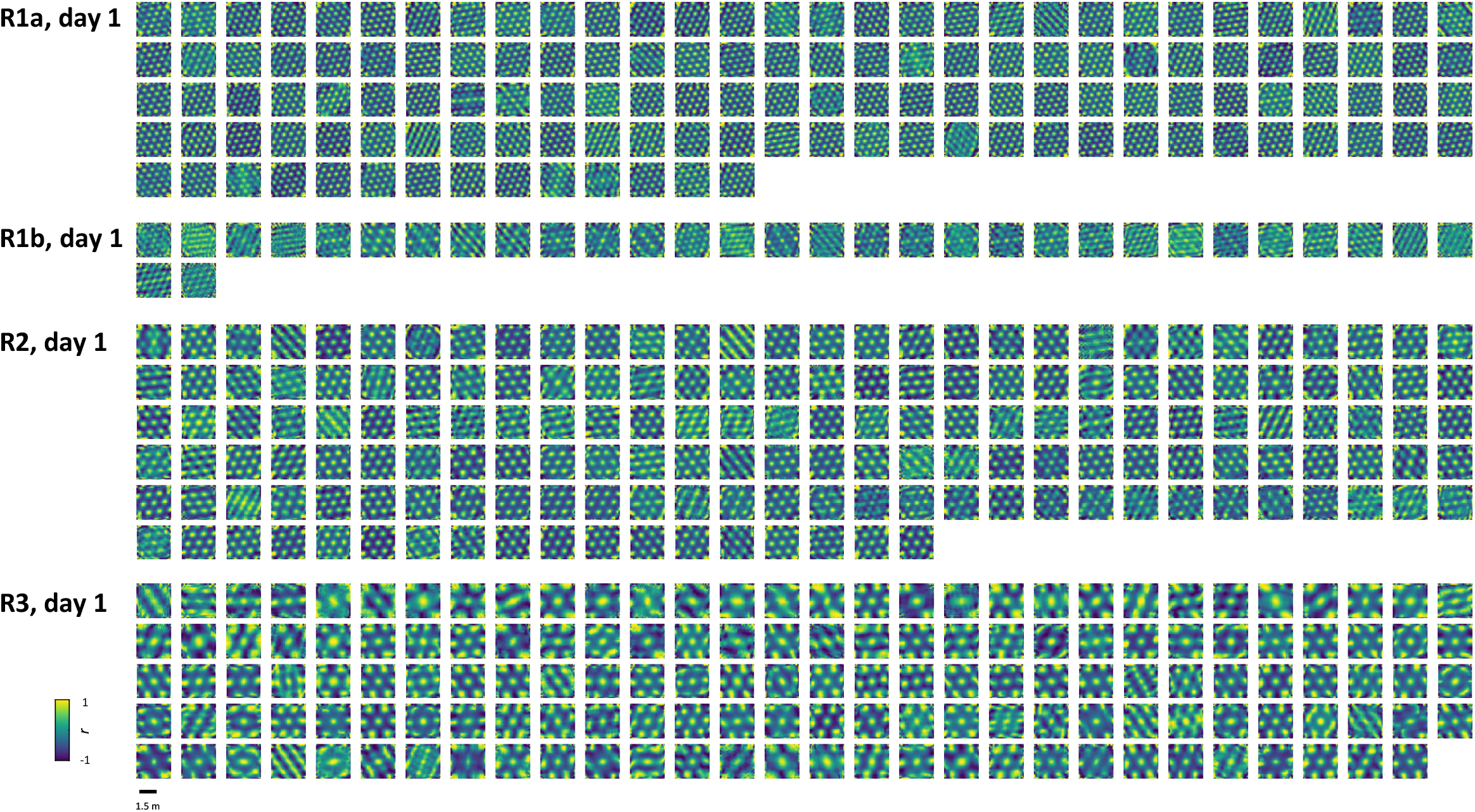
Spatial autocorrelograms of all grid cells. Same as Supplementary Figure 1 but showing data from rat R day 1 (total of 134 cells for module R1a, 32 cells R1b, 168 cells for R2, and 145 cells for R3).

**Supplementary Figure 3 for Reviewers:**
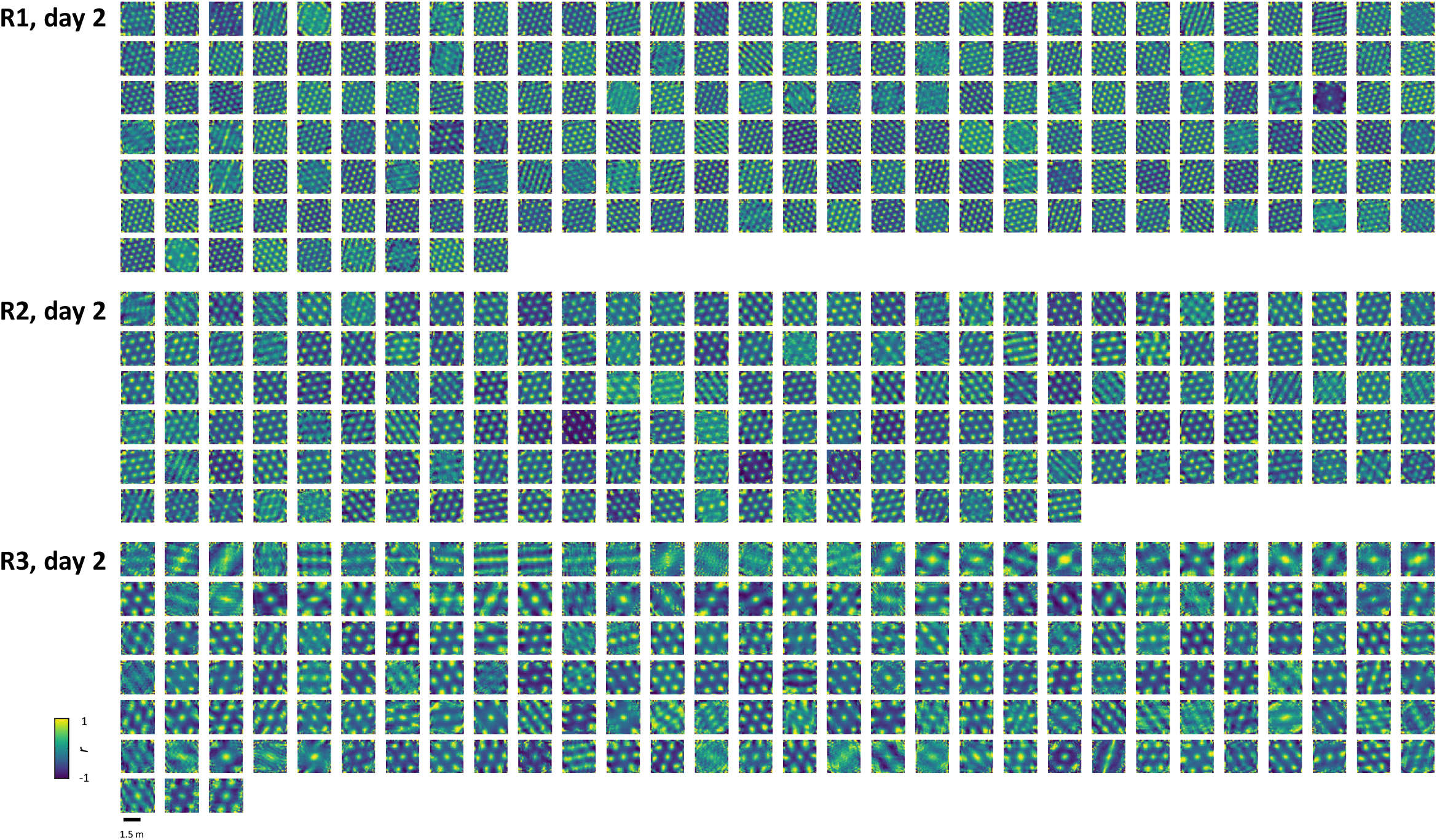
Spatial autocorrelograms of all grid cells. Same as Supplementary Figures 1 and 2 but showing data from rat R day 2 (total of 189 cells for module R1, 172 cells R2, and 183 cells for R3).

